# CD146^+^CD107a^+^ Mesenchymal Stem/Stromal Cells with Signature Attributes Correlate to Therapeutic Potency as “First Responders” to Injury and Inflammation

**DOI:** 10.1101/787176

**Authors:** Annie C. Bowles, Dimitrios Kouroupis, Melissa A. Willman, Carlotta Perucca Orfei, Ashutosh Agarwal, Diego Correa

## Abstract

CD146^+^ bone marrow–derived Mesenchymal Stem/Stromal Cells (BM-MSC) play key roles in the perivascular niche, skeletogenesis and hematopoietic support, however elucidation of therapeutic potency has yet to be determined. Here, inflammatory challenge to crude BM-MSC captured a baseline of signatures including enriched expression of CD146^+^ with CD107a^+^, CXCR4^+^, and LepR^+^, transcriptional profile, enhanced secretory capacity, robust secretome and immunomodulatory function with stimulated target immune cells. These responses were significantly more pronounced in CD146^+^ (POS)-selected subpopulation than in the CD146^-^ (NEG). Mechanistically, POS uniquely mediated robust immunosuppression while inducing significant frequencies of Naïve and Regulatory T cells *in vitro*. Moreover, POS promoted a pivotal M1-to-M2 macrophage shift *in vivo,* ameliorating inflammation/fibrosis of joint synovium and fat pad of the knee, failed by NEG. This study provides high-content evidence of CD146^+^CD107a^+^ BM-MSC, herein deemed ‘first responders’ to inflammation, as the underrepresented subpopulation within crude BM-MSC with innately higher secretory capacity and therapeutic potency.

**HIGHLIGHTS:**

- Signature phenotypic, transcriptional, and secretome profiles were identified and enriched in human CD146^+^ (POS)-selected subpopulation in response to inflammation
- Inflammatory challenge consistently altered stemness (*LIF*) and differentiation master regulators (*SOX9, RUNX2, PPARγ*) in crude, POS, and NEG BM-MSC, and deduced unique expressions in POS compared to NEG
- POS BM-MSC mediated the strongest immunomodulation, *e.g.* target immune cell suppression, Treg induction, diminished T cell differentiation
- POS BM-MSC promoted the largest M1-to-M2 shift *in vivo* alleviating induced synovitis and infrapatellar fat pad fibrosis of the knee

## INTRODUCTION

From the earliest studies that defined mesenchymal stem cells (MSC), the unique qualities inherent to these plastic-adherent cells still warrant detailed investigation into the dynamic responses induced by various molecular cues that influence their functional capacity, or potency. The *in vitro* multi-lineage differentiation capacity of MSC initially guided researchers to weigh heavily on engraftment and cell replacement strategies; however, we now recognize the innate paracrine signaling and immunomodulatory capacity of these cells as mechanisms to facilitate local changes in various milieus (Caplan and Correa, 2011; Salgado and Gimble, 2013; Salgado et al., 2018). The constitutively active functions of MSC are modulated by the surrounding environment, which informs MSC of spatial and temporal cues (Caplan, 2017a; Caplan and Sorrell, 2015). In response, MSC produce a tailored profile of secreted factors, miRNAs and extracellular vesicles (*i.e.,* collective secretome) instructive and conducive to the milieu leading to the understanding that the efficacy of MSC lie mainly in the combined potency of its secretome (Kehl et al., 2019; Salgado et al., 2015).

The malleable responsiveness of MSC has led researchers to directing these responses *in vitro* through cell pre-conditioning (*e.g.,* priming) (Kouroupis et al., 2018a). Subsequent interrogations of the cells and associated secretomes help correlate those responses to *in vivo* outcomes and to explore ways to enhance MSC potency (Tomchuck et al., 2008; Waterman et al., 2012; 2010). These dynamic features, including the immune cell-like responsiveness, secretory capacity, high migratory potential, and immunomodulatory cross-talk to other cells, along with their unique stem-like state and clonal reproducibility have made it difficult to distinguish the MSC beyond the minimum International Society of Cell and Gene Therapy (ISCT) criteria (Dominici et al., 2006; Krampera et al., 2013). Consequently, the once regarded “unique cell” dogma has changed, re-defining “mesenchymal stem cells” to “multipotent mesenchymal stromal cells” as stated by ISCT in 2005 (Horwitz et al., 2005) or “medicinal signaling cells” (Caplan, 2017b).

The perivascular prevalence of MSC in most tissues have led to the proposal of MSC and pericytes as developmentally related (Caplan, 2017a; da Silva Meirelles et al., 2009). Compelling evidence has demonstrated that MSC and pericytes do, in fact, share similar phenotypic markers (*e.g.,* CD146, NG2, PFGF-Rβ), the perivascular niche, differentiation potential and functional effects regarding tissue homeostasis and immunomodulation (Corselli et al., 2013b; Crisan et al., 2008; da Silva Meirelles et al., 2008; Gomes et al., 2018; Schwab and Gargett, 2007). Within the bone marrow (BM), CD146 expression helps to developmentally discriminate the bone and stroma from the cartilage component of BCSPs (Bone, Cartilage and Stroma Progenitors) arising from termed Skeletal Stem Cells (SSCs) (Chan et al., 2018), as well as perivascular from bone-lining/endosteal BM-MSC, all exhibiting osteogenic and hematopoietic support capabilities (Corselli et al., 2013a; Espagnolle et al., 2014; Tormin et al., 2011), and significantly affected by aging (Maijenburg et al., 2012). CD146^+^ BM-derived perivascular MSC are further discriminated based on Nestin and Leptin Receptor (LepR) expression (Mendelson and Frenette, 2014a; Méndez-Ferrer et al., 2010; B. O. Zhou et al., 2014), capable of forming heterotopic ossicles where they can transfer a hematopoietic functional tissue (Sacchetti et al., 2007). Finally, CD146^+^ pericytes, along with CD34^+^ adventitial cells within the perivascular space (Corselli et al., 2012) have been collectively called perivascular stem/stromal cells (PSC), suitable for bone reconstruction (James et al., 2017; 2012; Wang et al., 2019).

The significance of this semantic determination and the lack of specific markers to prospectively identify MSC leaves room for deep analysis of the function behind the phenotype that has yet to be greatly substantiated. By investigating the key pericyte marker CD146 on MSC, our objective is to not only shed light on the ambiguity of MSC phenotypes but provide compelling evidence of additional, less known signatures correlative to the unique immunomodulatory subpopulation of MSC with functions beyond its described osteogenic potential and hematopoietic support. The distinction is based on the hypothesis that a specific subset of MSC located within the perivascular niche (CD146^+^) would exhibit more robust responses to insults and inflammatory cascades than MSC otherwise more distant from vascular structures (CD146^-/low^). Such responses, in turn, greatly depend on the secretory capacity of the cells, intrinsically variable among populations and susceptible to be modulated by the immediate environment. Additional signature markers we describe herein as correlative with CD146^+^ BM-MSC are CD107a, CXCR4, and LepR. CD107a, or lysosomal-associated membrane protein-1 (LAMP-1), has been described as a marker of highly secretory cytotoxic T cells and actively degranulating natural killer (NK) cells (Alter et al., 2004; Kannan et al., 1996; Sudworth et al., 2016; Vego et al., 2016; Wattrang et al., 2015; York and Milush, 2015), but largely unknown in MSC. Thus, an assessment of the secretory status of the cells (*e.g.,* paracrine activity) would inform about how responsive they are at a particular time, susceptible to be associated with their potency. CXCR4 and its cognate ligand SDF-1/CXCL12 form an axis largely documented as mediating migrating responses of MSC to sites of injury (Kouroupis et al., 2018b; Liu et al., 2011). Leptin has been historically associated with energy regulation (Friedman and Halaas, 1998) and bone remodeling (Ducy et al., 2000), signaling through LepR in the hypothalamus and BM cells, respectively. In the BM, Leptin/LepR axis modulates osteogenic and adipogenic differentiation events of resident MSC (Scheller et al., 2010). Furthermore, LepR helps identify a multipotent progenitor population that are clonogenic in culture and give rise to osteoblasts and adipocytes in adult BM (B. O. Zhou et al., 2014). Morrison and colleagues further established that within those BM cells, LepR acts as a sensor of systemic energy inducing adipogenic while inhibiting osteogenic program in response to high fat diet (Yue et al., 2016). Collectively, LepR identifies BM progenitors with specific differentiation potential, susceptible to be modified by external factors.

A selection approach based on function and secretory status may help reduce the innate heterogeneity of isolated MSC, while characterizing a more uniform, robust, highly responsive cell subpopulation implicated behind the beneficial effects of a crude (*i.e.,* unfractionated) MSC preparation. Herein, we showed human BM-MSC (n=8) exhibited novel phenotypic (*e.g.,* CD107a^+^) and transcriptional signatures (*e.g., LIF*) after inflammatory priming unfractionated (UNF), correlated to the in-depth analysis of a higher secretory profile and reduced variability in CD146^+^ (POS) sorted cells compared with the CD146^-^ (NEG) fraction. Upon inflammatory challenge, secretion of numerous protein mediators at markedly higher levels from the POS subpopulation suggested a significantly more rapid response producing a robust immunomodulatory secretome, evident in reduced proliferation of stimulated target immune cells *in vitro* and M2 macrophages skewing *in vivo*. These *in vitro* responses were further translated using a rat model of acute inflammation/fibrosis of the joint synovial membrane and fat pad, where an intra-articular injection of POS cells consistently resulted in greater therapeutic efficacy compared with the marginal effects with the NEG fraction. Mechanistically, the therapeutic difference relates with a more M2-like (*i.e.,* alternative activation) macrophage phenotype occurred in POS-treated knees, indicative of anti-inflammation and repair mechanisms at the synovium and fat pad.

Together, this deep investigational approach into evaluating the MSC programs behind the pericyte marker CD146 provides compelling evidence that the CD146^+^CD107a^+^ BM-MSC subpopulation (correlative with CXCR4 and LepR expression) constitutes the therapeutically enhanced cells, or “first responders”, behind the crude MSC population, confounded by the heterogeneity captured during the unselective isolation process. Immediate implications relate with uniformity during cell-based product manufacturing, more reproducible outcomes and enhancing the precision for defining mechanistic underpinnings for bench-to-bedside translation.

## RESULTS

### Characterization of BM-MSC

BM-MSC from 8 donors of both sexes (36.5±9.8 years of age) (Figure S1A) were evaluated by standard characterization assays to validate BM-MSC qualities (Figure S1). Quantitative analysis of BM-MSC expressions included frequencies of positive surface markers CD105 (99.55±0.25%), CD90 (97.64±1.21%), CD73 (98.95±1.02%), and CD44 (98.26±1.42%), and negative surface markers CD34 (0.43±0.37%), CD31 (0.53±0.30%), CD45 (0.53±0.30%), CD31 (0.3±0.20%), and HLA-DR (0.58±0.36%; Figure S1C). Multipotency was assessed by trilineage differentiation assays. Quantification of Oil Red O, Alizarin Red, and glycosaminoglycans/alcian blue demonstrated BM-MSC capacity for induced adipogenesis (2.00±0.32 Normalized OD), osteogenesis (3.83±1.40 Normalized OD), and chondrogenesis (26.53±10.85; Figure S1D). Qualitative assessments showed morphological and growth kinetics indicative of BM-MSC, and further no observable differences when undergone inflammatory challenge/priming (Figure S1B and S1E). However, growth kinetics do show slight alterations in some donors (1, 4, and 6) depicting a response upon priming resulting in an initial delayed proliferation to iterate donor-specific differences that are ascertained by this priming method (Figure S1E).

### Signature profiles define human BM-MSC response to inflammation challenge

Phenotypic analysis of antigens designated as signature markers on the surface of the UNF Primed BM-MSC compared to the UNF Naïve cohort showed a markedly enhanced expression profile. Expression of marker CD146 was increased from 50.14±15.50% on UNF Naïve to 60.10±18.44% (*p<*0.05) on UNF Primed BM-MSC (Figure 1A). CD107a expression markedly increased from 9.70±6.07% on UNF Naïve to 43.68±21.43% (*p<*0.01) on UNF Primed BM-MSC, while CXCR4 increased from 11.50±9.72% on UNF Naïve to 38.47±17.55% (*p<*0.01) on UNF Primed BM-MSC. LepR expression increased from 5.09±1.97% to 28.55±15.28% (*p<*0.01) on UNF Naïve to UNF Primed BM-MSC, respectively (Figure 1B). Co-expression of primary signature marker CD146 with CD107a, CXCR4, or LepR led to an increased expression of 9.73±8.06% to 31.08±18.52% (*p<*0.01) for CD146^+^CD107a^+^, 10.97±8.69% to 32.05±11.17% (*p<*0.001) for CD146^+^CXCR4^+^, and 11.83±9.62% to 32.67± 17.40% (*p<*0.01) for CD146^+^LepR^+^ on UNF Naïve to UNF Primed BM-MSC, respectively, suggesting a signature phenotype consistently responding to inflammatory challenge in all donors (n=8; Figure 1C-D). Fluorescence images show the increased antigen presentation of each signature marker on UNF Naïve and Primed cohorts (Figure 1E and antibody controls in Figure S4A).

**Figure 1.**
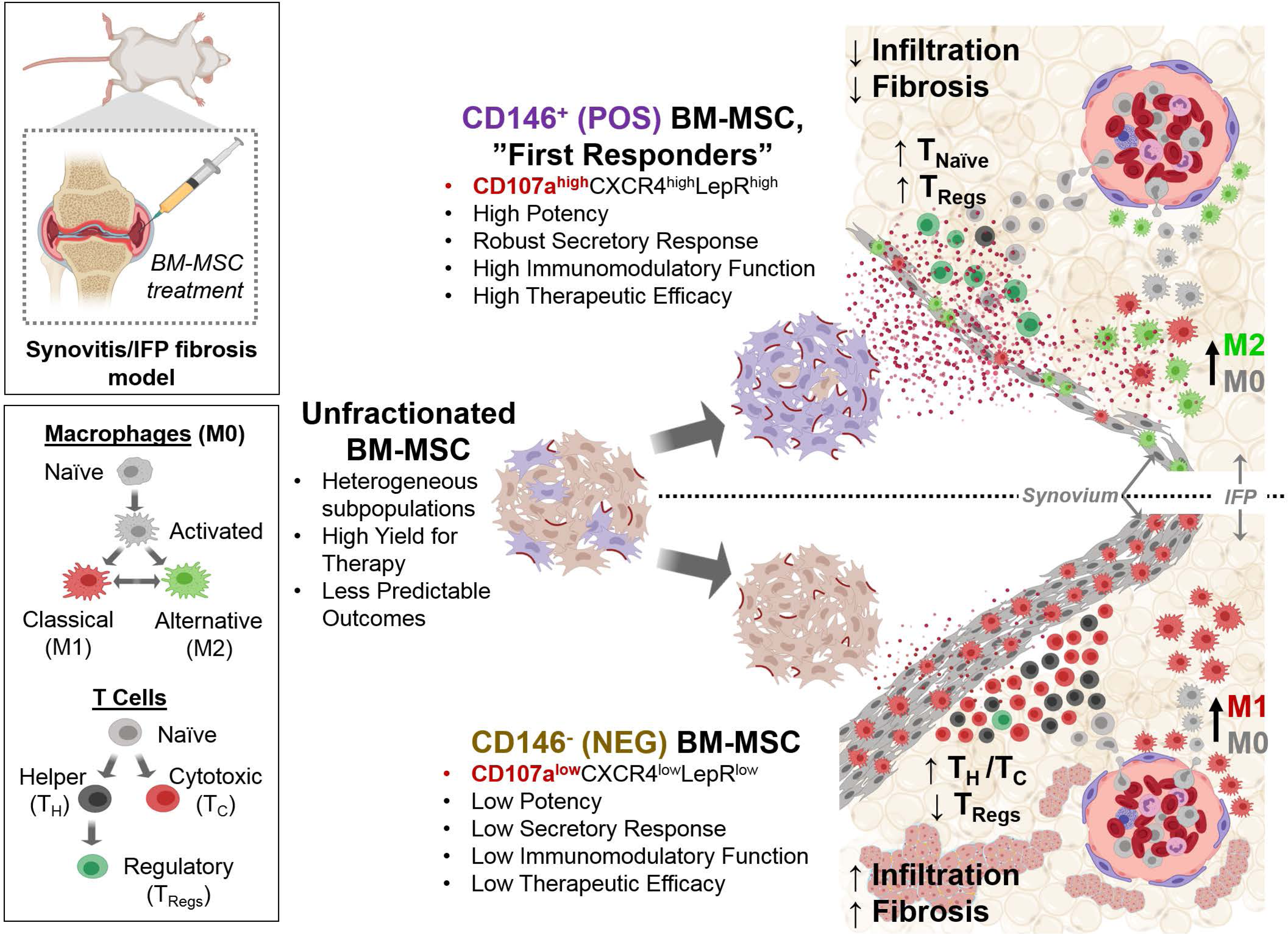
Unfractionated (UNF) BM-MSC expressed enrichment of defined signature markers upon inflammatory challenge (Primed). A. CD146 was designated as a highly distinguishable marker for a potently immunomodulatory BM-MSC subpopulation as extracellular expression was markedly increased in all UNF BM-MSC donors (n=8) when primed. B. Additional markers of interest used to distinguish a signature profile of immunomodulatory BM-MSC which expressions are comparably enhanced when challenged with inflammation. C-D. Images and quantitative analysis of the co-expressions of CD146 and designated immunomodulatory markers supported the signature markers highly enriched on a specific subpopulation of BM-MSC. E. Immunofluorescence of signature markers show constitutive expressions on UNF Naïve and increased expression in the UNF Primed. Scale bars represent 50 µm. Mean ± SD.

The marked enhanced CD146 expression and increased co-expression with signature markers CD107a, CXCR4, and LepR indicated that inflammatory challenge enriched a specific subpopulation within the crude UNF population. CXCR4 has been well established as an obligatory molecular marker of homing and migration, expressed on various cells including MSC. The inducible modulation of CXCR4 expression has led to many investigations establishing a focal stromal cell derived-1/CXCR4 axis that is able to facilitate therapeutic effects of BM-MSC during injury (M. Li et al., 2010; Liu et al., 2011; Wynn et al., 2004; Yang et al., 2015). Thus, the increased expression of CXCR4 during *in vitro* inflammatory challenge relates with a cell capable of mobilizing to injury sites. CD107a/LAMP1 (lysosome associated membrane protein-1) expression and functional implications on BM-MSC are largely unknown, until now. CD107a expressed intracellularly on endosomal membranes have been reported on immune cells, most notably cytotoxic T cell and natural killer cells, as indicators of activation or degranulation, respectively (Alter et al., 2004; Kannan et al., 1996; Sudworth et al., 2016; Vego et al., 2016; Wattrang et al., 2015; York and Milush, 2015). Likewise, our data suggests a similar CD107a expression correlative with an activated and secretory function after inflammatory challenge. Given that MSC express Toll-like receptors that are responsive to external stimuli (Tomchuck et al., 2008; Waterman et al., 2010), as do immune cells, increased CD107a expression now can be implicated as a functional signature of actively secreting BM-MSC. LepR expression on BM-MSC was previously reported as a perivascular cell used to track migration in the periarteriolar to perisinusoidal niche in response to hematopoietic stem cell cues (Mendelson and Frenette, 2014b). Moreover, LepR^+^ cells were found on perivascular, stromal populations with self-renewal capacity, suggested as source of osteoblasts and adipocytes in the adult bone marrow (Niu et al., 2015; Yue et al., 2016; B. O. Zhou et al., 2014). An additional study described a link to LepR expression and advanced age (Laschober et al., 2009), however here, LepR expression demonstrated inducible plasticity when exposed to an inflammatory challenge along with CD146, reminiscent of the perisinusoidal BM-MSC (Mendelson and Frenette, 2014a; Méndez-Ferrer et al., 2010). The parallel CXCR4 and CD107a enrichment suggests a transiently enriched active state expressed on a specific subpopulation of BM-MSC with high injury sensitivity and secretory capacity.

### Inflammatory challenge induced an immunomodulatory BM-MSC defining secretory and transcript profiles with indicators of high potency

Numerous growth factors and inflammation-related proteins were quantitatively compared between UNF Naïve and Primed cohorts, represented as heat maps to show relative fold changes (Figure 2A). Corroboration of the significantly altered proteins of the UNF Primed profile with pathway analysis further supported robust immunomodulatory signaling with the top 3 relevant pathways matched as IL-10 (HSA-6783783), Toll-like receptor (hsa04620), and IL-4 and IL-13 (HSA-6785807) signaling pathways (Figure 2B). Immunomodulatory activities facilitated by prostaglandin E-2 (PGE-2) and IDO were suggested to strongly influence the functional effects of the UNF Primed BM-MSC which were measured significantly higher (3.73±3.03-fold and 5.12±3.39-fold) compared to UNF Naïve cohorts, respectively (Figure 2C).

**Figure 2.**
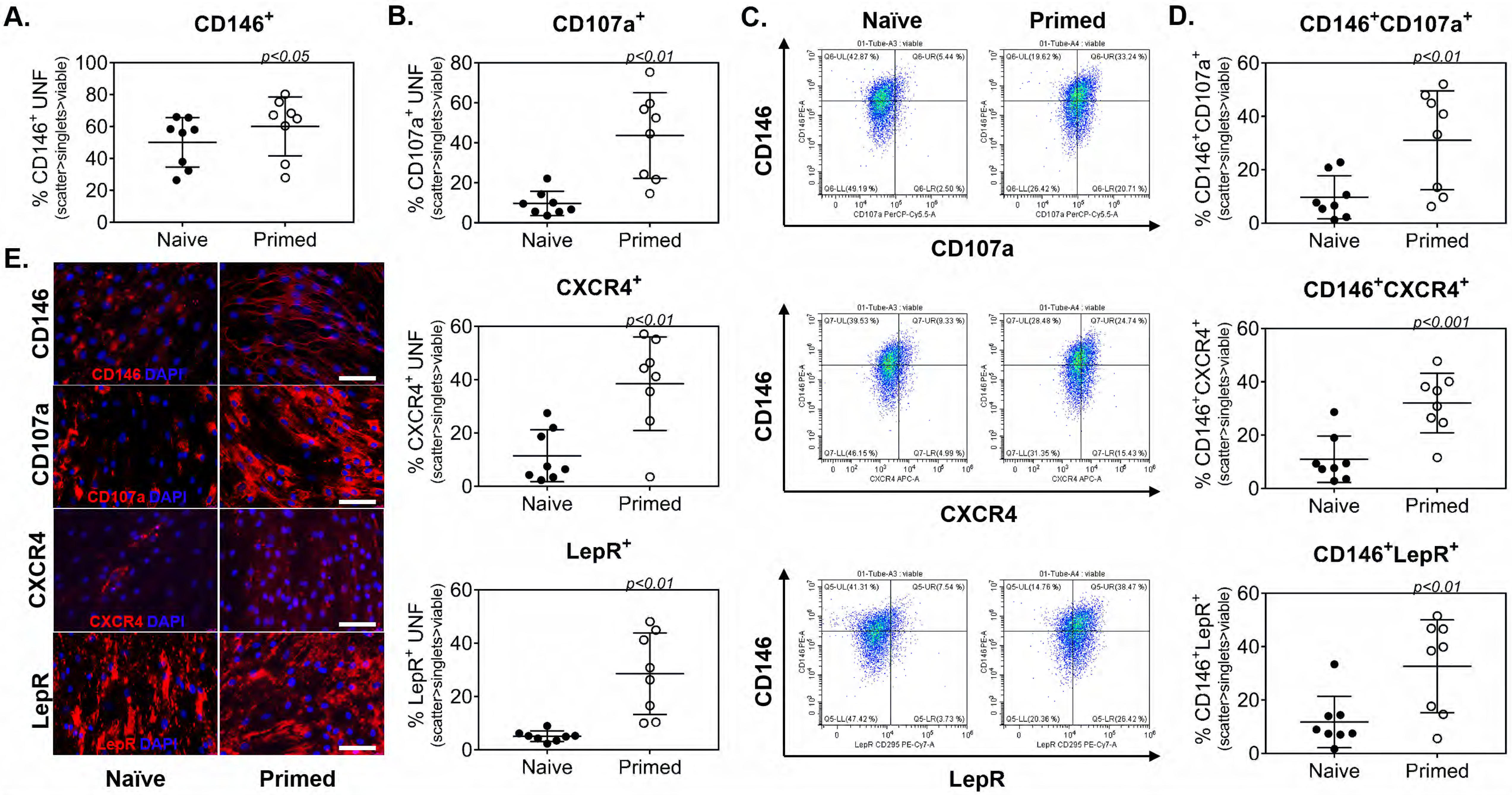
Immunopotency assay demonstrated greater potency of UNF Primed group. A. Quantitative analysis of proliferation rates indicated that UNF Primed BM-MSC diminished proliferation of stimulated T cells compared to UNF Naïve. B. Gating strategy for detection and phenotypic analysis of stimulated CFSE^+^ T cells. Phenotypic analysis of T cell subsets resulting from the direct co-culture of stimulated T cells with UNF Naïve and UNF Primed BM-MSC. D. Time-lapse images of every 24 hours during the 72-hour immunopotency assay visualized the CFSE^+^ T cells (green) co-cultured on top of the adherent UNF Naïve or Primed BM-MSC groups. Scale bar represents 300 µm. Mean ± SD. *, *p*<0.05; **, *p*<0.01; ***, *p<*0.001; ****, *p<*0.0001.

Similarly, the transcription profiles of genes revealed a consistent shift in transcripts in the UNF Primed BM-MSC compared to the UNF Naïve cohort induced by the inflammatory challenge. Several over-expressed and under-expressed genes were reproducibly and significantly up- or down-regulated, respectively, in all donors using the priming method (Figures 2D and S5A). Of these, UNF Primed BM-MSC showed up-regulation of leukemia inhibitory factor (*LIF*; 3.92-fold; *p*<0.0013), indicative of a highly immunosuppressive/tolerogenic immunity state and stemness (Metcalfe et al., 2015; Nasef et al., 2008). *LIF* has been related with Tregs enrichment and with an antagonistic effect on the pro-inflammatory cytokine IL-6 (Gao et al., 2009; Metcalfe, 2011), critical for the functional effects in immunomodulation (below). On the contrary, the three master regulators of mesenchymal differentiation programs were significantly down-regulated, namely *RUNX2* for osteogenesis (-4.11-fold; *p*<0.0012), *SOX9* for chondrogenesis (-2.91-fold; *p*<0.012), and *PPARγ* for adipogenesis (-2.49-fold; *p*<0.048) (Figure 3D). Interestingly, various other transcripts associated with those programs (e.g., *COL1A1, GFD5, BMP6, BGLAP*, *TGFβ1-3, ITGB1* etc.), were similarly downregulated (Figures 2D and S5A). These findings may suggest a potential dynamic phenotypic switch of the cells between progenitor and immunomodulatory capabilities, depending on the environment they see. Further investigation is warranted to clarify this potential phenotypic adaptability. Nevertheless, this evidence supports the use of inflammatory challenge to induce a robust signature phenotype of BM-MSC with potent immunomodulatory and stem-like gene expression and secretion distinct from its Naïve cohort.

**Figure 3.**
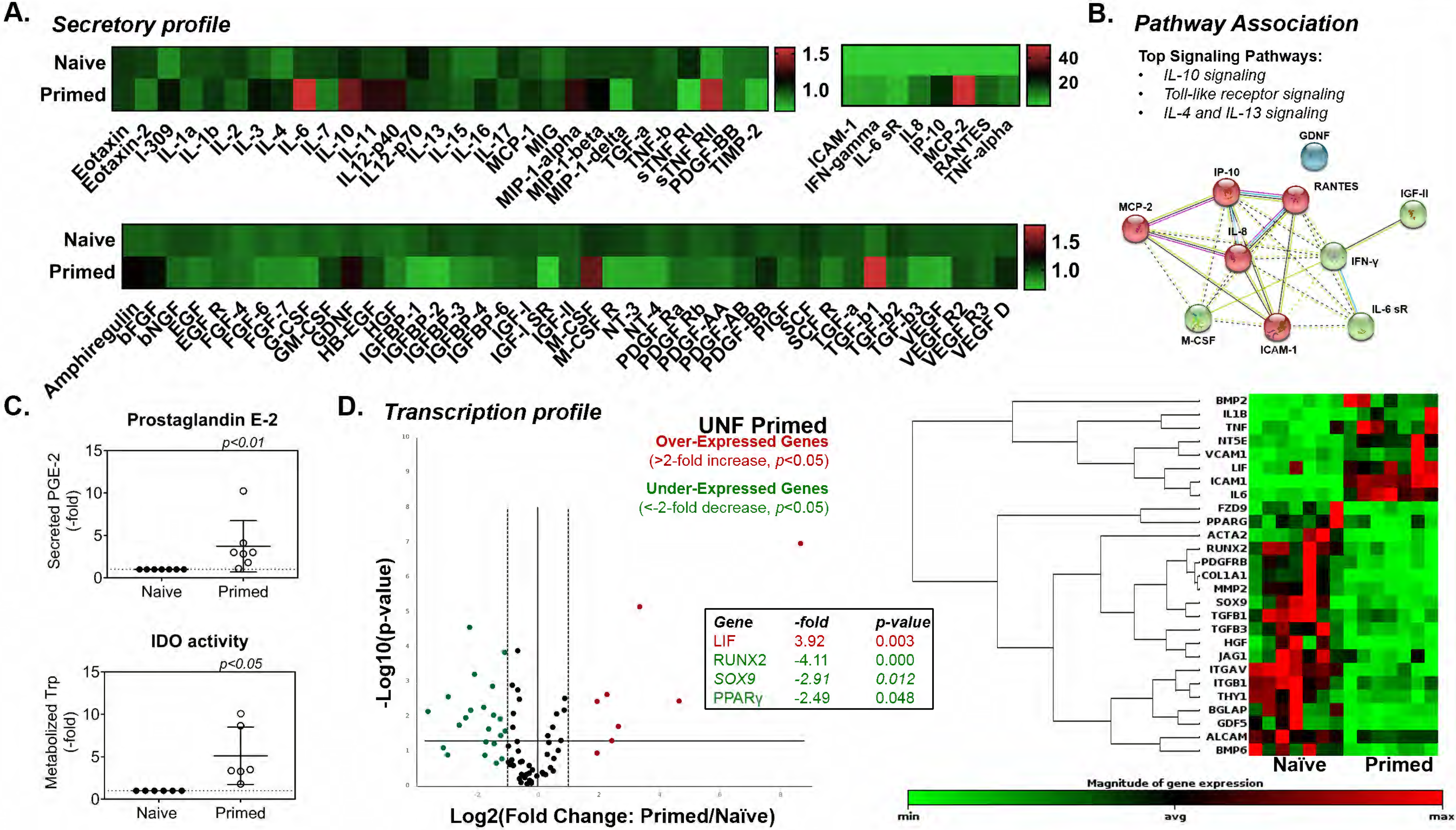
Secretory and transcriptional analyses showed distinguishable differences between UNF Naïve and UNF Primed BM-MSC. A. Enzyme-linked immunosorbent assays detected and revealed relative fold changes in the profile of secreted proteins of UNF Primed BM-MSC compared to UNF Naïve cohorts. B. Immunomodulatory proteins prostaglandin E-2 (n=6) and indoleamine 2,3-dioxygenase (IDO; n=7) were measured at markedly greater levels in UNF Primed BM-MSC than UNF Naïve cohorts. C. Analysis of transcription in UNF Primed cohorts compared to UNF Naïve showed significantly increased transcripts of stem-like and immunomodulatory products and significantly decreased transcripts that commit BM-MSC to multiple differentiated states (n=6). D. Unsupervised hierarchical cluster diagram of genes expressions that were over-expressed (>2 fold) and under-expressed (<-2 fold) in each group (n=6). Mean ± SD. *, *p*<0.05; **, *p*<0.01; ***, *p<*0.001; ****, *p<*0.0001.

### Immunomodulatory potency of UNF Primed BM-MSC mitigated PBMC and T cell proliferation and differentiation

The ability to assay the therapeutic potency of MSC *in vitro* is a context-dependent endeavor given the dynamic, robust, comprehensive and variable nature of MSC. Several molecules such as interleukin-10 (Jiao et al., 2011), prostaglandin E-2 (Solchaga and Zale, 2012), and IDO (Tomchuck et al., 2008; Waterman et al., 2010) have been examined to suggest potency of MSC, but given the robust secretory repertoire of MSC, a singular molecular assessment is inadequate. In this study and several others (Chinnadurai et al., 2018; Jiao et al., 2011), potency corresponded to diminished proliferation of stimulated immune cells when co-cultured with BM-MSC (Chinnadurai et al., 2018; Scruggs et al., 2013), further analyzing specific subsets of PBMC and T cells to determine immunomodulatory potency yielded more compelling implications.

According to the immunopotency assay (IPA) matrix (Figure S2), a dose-response evaluation of PBMC stimulated with PMA/ionomycin cocktail for a robust proliferative response was performed to distinguish the ratios and suppressive effects of UNF Naïve and UNF Primed BM-MSC. This evidence indicated that a 2:1 PBMC to BM-MSC ratio was optimal for achieving a significantly attenuated proliferative response of stimulated PBMC (No BM-MSC; 37.50±9.98%) by co-culture with UNF Naïve (21.00±19.61%; *p*<0.01) and, more so, UNF Primed BM-MSC (12.25±11.62%; *p*<0.001; Figure 3A). Higher ratios of PBMC to UNF Naïve BM-MSC progressively reduced the effects, measured at 4:1 (43.25±21.00%), 12:1 (76.75±10.75%), and 60:1 (78.00±10.92%). UNF Primed cohorts, on the other hand, produced a stronger immunosuppressive response at 4:1 (4.75±1.71%; *p*<0.0001), 12:1 (22.75±30.30%; *p*<0.05), and 60:1 (72.50±8.10%) ratios, overcoming the dose-dependent reducing effect observed with UNF Naïve BM-MSC.

To complement the data obtained with PBMC, human pan T cells were co-cultured (2:1) with same BM-MSC groups. T cells were stimulated with CD3/CD28 antibody cocktail which mimics a physiological activation by antigen-presenting cells (Trickett and Kwan, 2003) that warrants a minimal proliferative response over the 3-day IPA according to the experimental design (Figure S2). Control T cells (No BM-MSC) had a proliferation rate of 11.56±8.86%, significantly increased with co-culture with UNF Naïve BM-MSC (32.11±16.85%, *p*<0.05) but not with UNF Primed cohorts (20.30±17.62%) (Figures 3B and S4B), suggesting an immunosuppressive effect of the latter. Given the co-culture cell ratios used, the presence of BM-MSC were expected to increase T cell proliferation (Y. Zhou et al., 2013). However, a more detailed description of the proliferating cells was visualized as scatter patterns gated as parent T cells (CFSE^high^) and progeny T cells (CFSE^low^) using flow cytometric analysis for quantification of proliferation rates and phenotypic subsets (Figure 3C). Interestingly, despite the increased number of total proliferating T cells described (Figure 3B), the number of progenies visualized was reduced with UNF Naïve and more so with UNF Primed BM-MSC, suggesting reduced numbers of actual cell divisions during the co-culture (Figure 3C). Furthermore, and as presented later, the contributions of specific subpopulations of cells within the UNF BM-MSC help explain this phenomenon.

Stimulated T cells co-cultured with UNF Naïve (13.18±3.83%; *p*<0.001) and UNF Primed (12.82±1.65%; *p*<0.01) BM-MSC were significantly higher in frequency of CD3^+^CD4^-^CD8^-^ Naïve T cells than controls (7.16±3.14%), suggesting mitigated differentiation by both UNF BM-MSC cohorts rendering T cells in their naïve state (Figures 3D and S4C). The decreased trend of cytotoxic T cells resulting from the IPA with UNF Naïve (27.24±9.70%) and UNF Primed (26.50±3.73%) compared to No-BM-MSC controls (30.35±17.03%) suggested that BM-MSC did not promote differentiation of CD3^+^CD8^+^ Cytotoxic T cells. IPA results showed UNF Primed BM-MSC substantially reduced the differentiation to CD3^+^CD4^+^ Helper T cells to 45.50±9.00% (*p*<0.05) compared to controls (60.78±12.88%) while UNF Naïve cohorts (48.65±7.66%) slightly mitigated Helper T cell differentiation. Moreover, the percent of regulatory T cell (Treg) subsets (CD3^+^CD4^+^CD25^+^foxp3^+^) induced within the Helper T cell population were markedly enhanced by the co-culture with UNF Naïve (28.48±19.45%; *p<*0.05) and UNF Primed (25.36±22.63%) BM-MSC compared to No BM-MSC controls (5.56±3.87%; Figures 3D and S4C). Time-lapse analysis further demonstrated the proliferation responses of each T cell group during the IPA by CFSE detection (Figure 3E). Together, these data suggest that despite the increased number of T cells observed when BM-MSC are present, those T cells enlarge the pool of undifferentiated (i.e., naïve) cells and skew the Helper T cells towards an immunomodulatory Treg phenotype.

### POS BM-MSC express an enrichment of signature markers

CD146 was the main signature marker used to capture the largest subpopulation of BM-MSC responsive to an inflammatory challenge and is suggested to correspond with the immunomodulatory potency along with additional signatures also enhanced by priming. To further substantiate the immunomodulatory potency of BM-MSC related directly to expression of CD146, magnet-activated cells sorting (MACS) was used to fractionate UNF BM-MSC (n=3) into CD146+ (POS) and CD146- (NEG) fractions, which were subsequently interrogated by similar approaches performed on UNF BM-MSC. CD146-based MACS yielded POS (71.01±7.03%) and NEG (18.82±5.69%; *p*<0.05) subpopulations with marked differences in CD146 expression, although sub-optimal for capturing pure subpopulations (Figure 4A). This might be explained by the reported non-discrete expression of CD146 in BM-MSC, exhibiting high/bright, low/dim expressivity levels (Espagnolle et al., 2014), captured differently by the MACS-associated antibody and the one used for phenotypic characterization.

**Figure 4.**
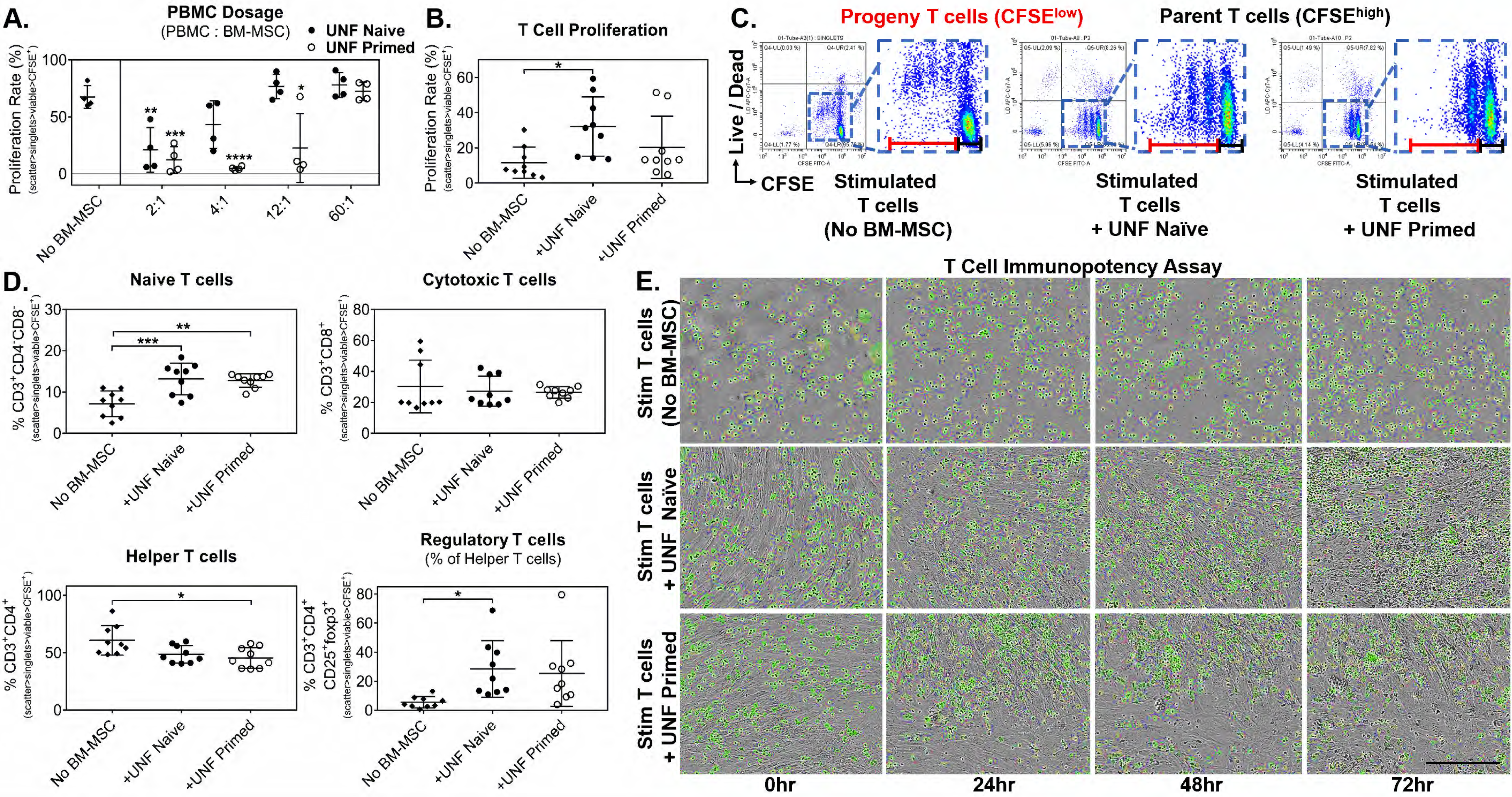
Sorting by CD146 allowed the identification of inherent features associated with CD146^+^ (POS) and CD146- (NEG) BM-MSC. A. Magnet-activated cell sorting (MACS) was used to capture CD146^+^ (POS) and CD146- (NEG) BM-MSC. B. Signature marker were differentially expressed and enriched in Naïve and Primed cohorts of POS and NEG BM-MSC (n=8). C. Secretory potency was measured as total secreted molecules and represented as time-lapse secretion as well as the area under curve for the 48-hour duration of priming (n=4). C. Immunofluorescence of signature markers on the extracellular membranes of POS and NEG BM-MSC. Scale bars represent 50 µm. Mean ± SD. *, *p*<0.05; **, *p*<0.01; ***, *p<*0.001; ****, *p<*0.0001.

Phenotypic analysis corresponding to Naïve and Primed cohorts of POS and NEG BM-MSC subpopulations supported the enrichment of signature markers when exposed to inflammatory challenges similar to those evidenced in the UNF BM-MSC population. CD146 expression was detected in the POS Naïve (84.45±6.11%) and NEG Naïve (16.08±6.12%) cohorts which was further enriched with priming in both the POS Primed (91.9±6.49%; *p*<0.05) and NEG Primed (23.44±8.93%) cohorts, suggesting a permanent responsiveness of the cells independent of the basal CD146 expression level. Highly significant differences remained between the POS Naïve and NEG Naïve (p<0.0001) and the POS Primed and NEG Primed (*p*<0.0001) cohorts.

Expression of CD107a was significantly increased in both the POS and NEG Primed BM-MSC (36.51±17.76% *p*<0.05 and 21.77±13.21%; *p*<0.05, respectively), compared to the corresponding POS and NEG Naïve cohort (15.09±9.64% and 8.87±6.03%), respectively, maintaining a higher expression in the POS group overall. Similarly, CXCR4 and LepR expressions were increased with priming in both the POS and NEG BM-MSC subpopulations. POS Naïve BM-MSC (15.31±12.24%) expression of CXCR4 increased to 35.70±20.38% on POS Primed cohorts while NEG Naïve (8.85±7.28%) was increased to 24.11±16.85% on NEG Primed cohorts. Modest enrichment of LepR was measure from 13.16±15.97 expression on the POS Naïve to 30.18±27.29% on POS Primed and remained comparable between NEG Naïve (6.33±8.04%) and NEG Primed (12.19±11.88%) BM-MSC (Figure 4B). As with CD107a, CXCR4 and LepR showed overall higher, yet not statistically significant, levels in the POS group. Fluorescence images of POS and NEG BM-MSC validated the increased expression of signature markers as Naïve BM-MSC compared to Primed cohorts at 48 hours of inflammatory challenge (Figure 4C). Together, these phenotypes verify that the POS subpopulation was enriched with the signature profile designated as a distinguishable subset of BM-MSC that produce a markedly higher response to inflammation *in vitro*.

### Secretory potency and profile of POS BM-MSC was most robust and immunomodulatory

By staining BM-MSC groups with CFSE, secretory potency was measured as total extracellular release of intracellular contents by Primed BM-MSC normalized to Naïve cohorts throughout the 48-hour inflammatory challenge. Initially, both POS Primed and NEG Primed BM-MSC responded in a large increase of released contents captured within the first hour then tapered off by hour 4, which is likely attributed to the media change. Consistently, POS Primed BM-MSC released a higher total of contents throughout the detected time points of the 48-hour priming period. Secretory release by hour 4 had equilibrated the cells to constitutive release measured by POS Primed (1.57-fold) and NEG Primed (1.36-fold) and remained steady until a final drop to comparable levels of the Naïve cohorts at 48 hours (Figure 4D, top). To further validate the larger potency of the POS subpopulation compared to the NEG subpopulation, the cumulative secretory potency of each BM-MSC group was quantitatively compared. POS Naïve (59.38±11.65) and POS Primed (96.58±32.88; *p*<0.05) groups generated a more robust secretory release over 48 hours than released by NEG Naïve (41.38±5.35) and NEG primed (58.83±11.97) groups (Figure 4C, bottom). In fact, the constitutive secretory release of contents of the POS Naïve group was indistinguishable to the amount released by the NEG Primed cohort over the 48 hours demonstrating the greater secretory potency of the POS subpopulation over the NEG subpopulation. Interestingly, this secretory profile mirrors the expression profile of CD107a in all groups, stressing the correlation between phenotype and function.

To elucidate the molecules that contribute to the compelling secretory data distinguishable of the POS subpopulation, analyses of immunomodulatory PGE-2 and IDO activity were measured to substantiate potency. Constitutive IDO activity, indirectly measured by metabolized tryptophan (Trp), NEG Naïve BM-MSC was measured as 0.47±0.26 pmole (*p*<0.05) which was significantly less than the activity measured by POS Naïve (1.90±1.16 pmole). As reported by UNF BM-MSC, inflammatory challenge further enhanced IDO activity with POS Primed increasing to 2.59±2.03 pmole and NEG Primed to 2.12±2.3 pmole (Figure 4F).

Over 80 proteins were detected and generated into secretory profiles for the POS and NEG BM-MSC groups throughout the 48-hour inflammatory challenge. Heat map representation of the relative fold changes between the Primed and Naïve cohorts of the UNF, POS, and NEG BM-MSC demonstrated the largest increases amongst the inflammation-related growth factors and cytokines corresponding to the POS Primed group (Figure 5A-B). Moreover, quantitative comparisons were presented individually of pertinent immunomodulatory proteins secreted by each Primed cohort normalized to the Naïve cohort of the UNF, POS, or NEG BM-MSC. Of the most significant proteins, POS Primed secreted the highest levels of transforming growth factor-β1 (TGF-β1; 3.22-fold; *p*<0.0001), soluble tumor necrosis factor receptor 2 (sTNF-R2; 2.61-fold; *p*<0.01), macrophage inflammatory protein-1α (MIP-1α; 4.33-fold; *p*<0.01), platelet derived growth factor-BB (PDGF-BB; 1.46-fold; *p*<0.05), interleukin-10 (IL-10; 1.84-fold; *p*<0.05), interleukin-12 p40 (IL-12 p40; 1.57-fold; *p*<0.05); interleukin-13 (IL-13; 1.88-fold; *p*<0.05), interleukin-15 (IL-15; 1.73-fold; *p*<0.05), and macrophage-colony stimulating factor (M-CSF; 1.31-fold; *p*<0.05). Macrophage chemotactic protein-2 (MCP-2) and Regulated upon Activation, Normal T cell Expressed, and Secreted (RANTES) were both significantly greater in the supernatants of the UNF Primed measured as 84.71-fold (*p*<0.0001) and 28.75-fold (*p*<0.05) and POS Primed measured as 54.05-fold (*p*<0.001) and 19.49-fold (*p*<0.05), respectively. Interestingly, interleukin-6 soluble receptor (IL-6sR; 4.79-fold; *p*<0.05) and inflammatory protein-10 (38.07-fold; *p*<0.01) were most significant secreted in the UNF Primed compared to the individual subpopulations suggesting synergistic effects captured in UNF BM-MSC from the POS and NEG subpopulations. Nevertheless, the POS Primed BM-MSC had greater levels of IL-6sR and IP-10 measured as 2.82-fold and 20.72-fold compared to the NEG Primed levels measured as 2.02-fold and 14.47-fold, respectively. Inflammation challenge induced the largest secretory increases of intracellular adhesion molecule-1 (ICAM-1), interleukin-8 (IL-8), and interferon-γ (IFN-γ) in all BM-MSC groups which are suggested to be directly correlated with exogenous IFN-γ and TNF-α use in the inflammatory challenge. ICAM-1 was highest in the supernatant of POS Primed groups (9.20-fold; *p*<0.001) followed by NEG Primed (8.14-fold; *p*<0.01) and UNF Primed (6.92-fold; *p*<0.05) while IL-8 was highest in the UNF Primed groups (9.74-fold; *p*<0.001) followed by NEG Primed (8.16-fold; *p*<0.01) and POS Primed (7.54-fold; *p*<0.05). Although detection of IFN-γ included secreted and exogenous protein used for priming, levels were comparable in the UNF Primed (11.31-fold; *p*<0.001), POS Primed (10.19-fold; *p*<0.001), and NEG Primed (9.56-fold; *p*<0.001) BM-MSC (Figure 5C).

**Figure 5.**
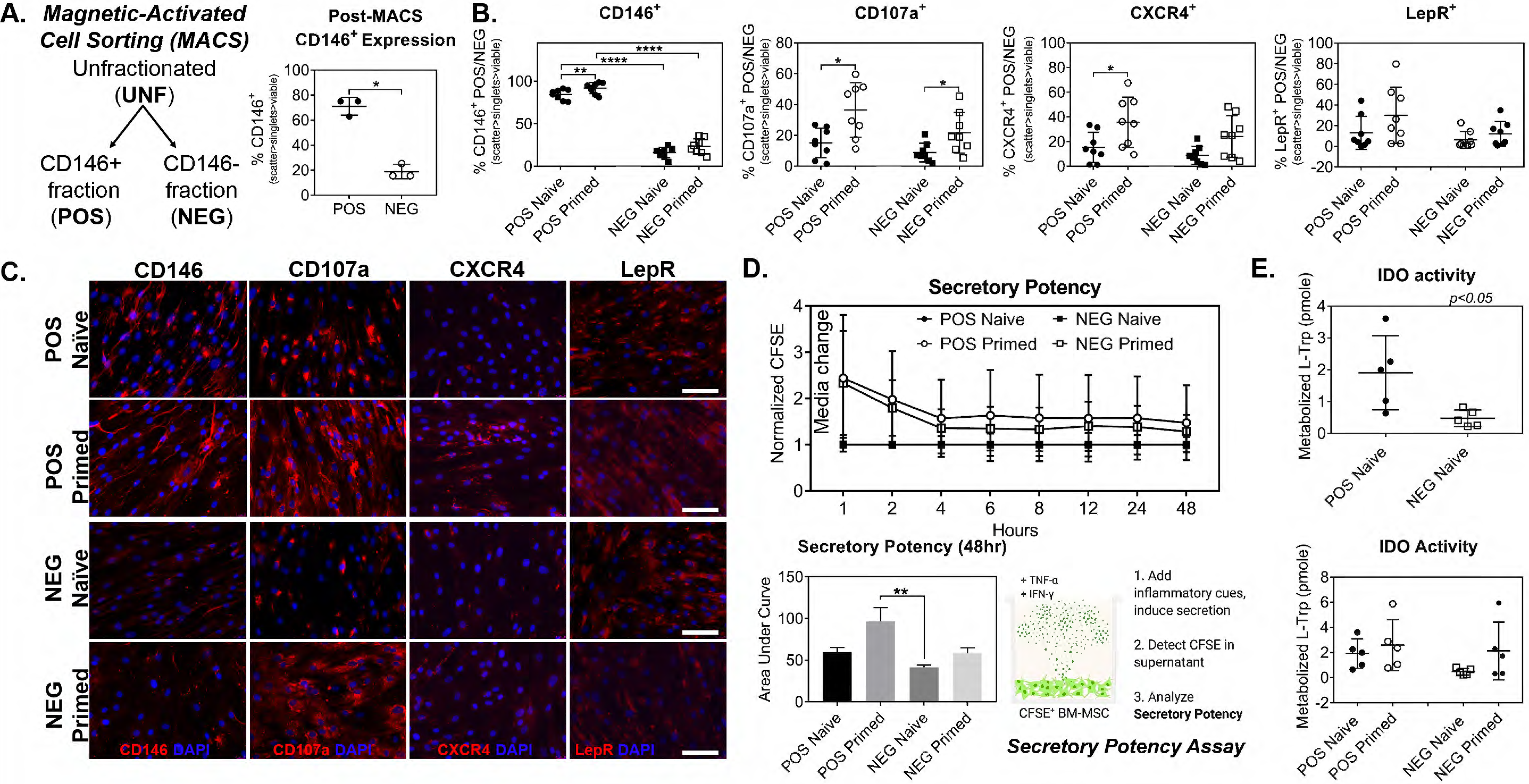
Secretory profiles support that the POS BM-MSC are the potent responders to inflammation. A-B. Fold change differences between the Naïve and Primed cohorts of the UNF, POS and NEG groups (n=8) reveal profiles indicating the largest secretory profiles of inflammatory and growth factor mediators by the POS BM-MSC with priming. C. Protein mediators determined to have the most significant differences amongst each group, highlighting the most robust changes of immunomodulatory factors by the POS BM-MSC.

Of the 16 proteins discussed that were measured as significantly greater from at least one of the Primed cohorts of UNF, POS, or NEG BM-MSC, 14 were from the POS BM-MSC subpopulation, supporting their robust and comprehensive secretory profile in response to inflammatory challenge. Pathway associations with all significantly altered proteins reveal differences in the signaling pathways between subpopulations, where Rheumatoid Arthritis (RA)-associated (hsa05323), Jak-STAT (hsa04630), TNF (hsa04668) and IL-17 (hsa04657) pathways were the top matches as modulated by the POS subpopulation (not the NEG) (Figure 6A, top). All 3 groups shared only 4 altered factors. Furthermore, unlike with UNF (2) and NEG (2), several of the altered factors (10) were exclusively altered in the POS group. POS shared more with the UNF (6) than with NEG (3), while NEG did not coincide with UNF in any response. This suggests a selective contribution of the POS within crude preparations to the immunomodulatory signaling for mechanistic targets and insight, especially related with inflammation and autoimmunity. Radar profile graphs from categorized signaling activities show high similarities and alignment between the UNF and POS subpopulation in both Biological Process and Reactome, especially with effects on leukocyte proliferation (GO:0070663) and TNF signaling (hsa04668) (Figure 6A, middle and bottom). These data further bolster that it is the effects of the POS subpopulation that heavily influences the overall effects detected by the UNF BM-MSC.

**Figure 6.**
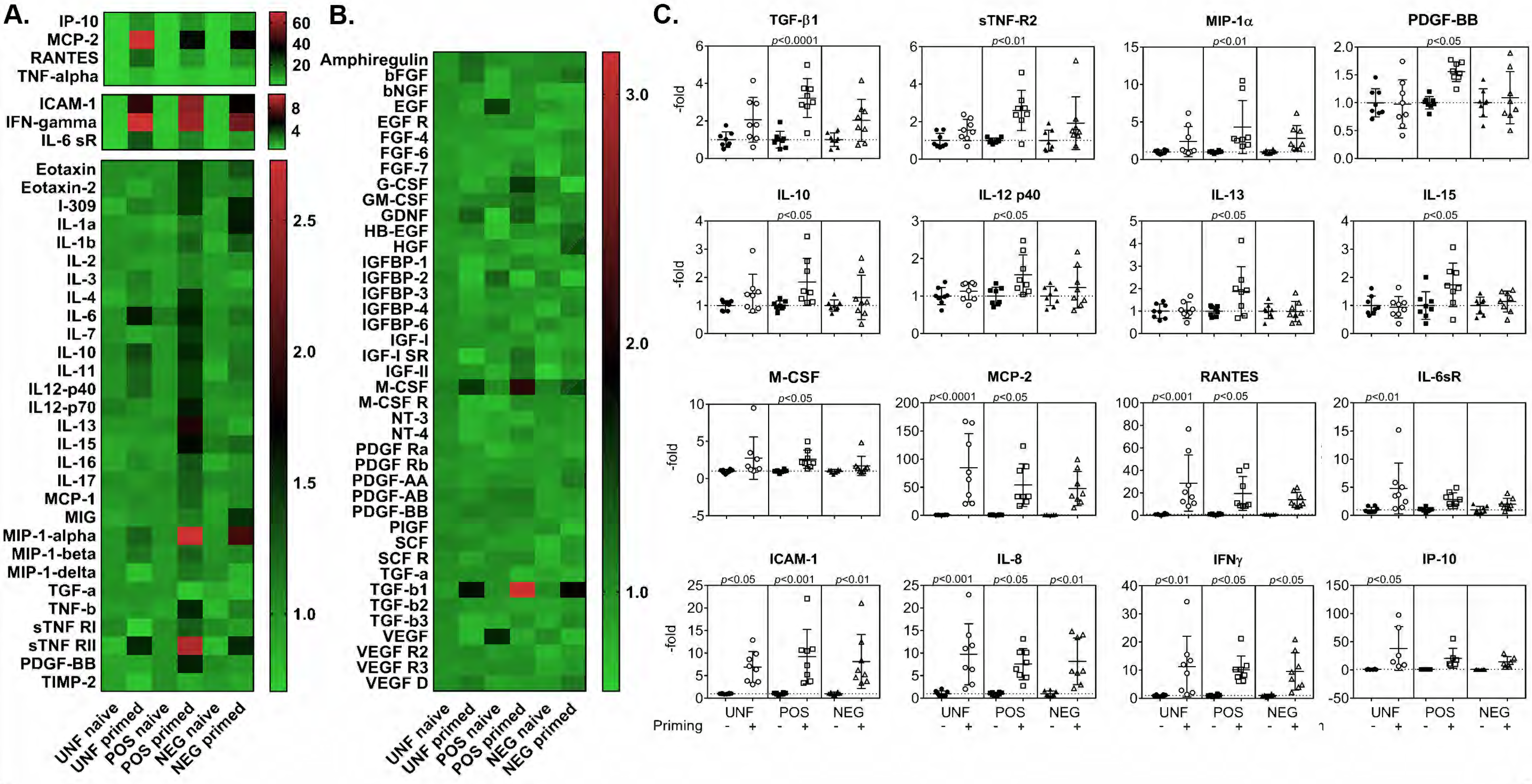
Pathway analysis of secreted proteins and transcriptional profiles. A. Highly regulated proteins (>1.4-fold increase and <-1.4-fold decrease) were oriented with STRING to derive the involvement of relevant pathways. Venn diagram, Biological Process, and Reactome associates exhibit the distinctive properties and the shared overlaps among the UNF, POS, and NEG BM-MSC. B. Transcriptional profiles of the POS and NEG subpopulations in response to inflammatory challenge show distinguishable difference leading to identification of key transcripts that are key associations with the specific subpopulation. Venn diagram further displays the strong functional influence of the POS subpopulation underlying the UNF BM-MSC. Stars represent statistical significance with values greater than 2-fold or less than -2-fold change.

### Transcription profiles reveal signatures distinguishable of both subpopulations

Comprehensive strategies to determine the influence of the POS or NEG subpopulations within the cumulative UNF BM-MSC are necessary to understand the contribution of each that may guide investigations to reducing heterogeneity for more consistent outcomes. The transcription profiles compared the genes actively over- or under-expressed following inflammatory challenge in each BM-MSC group (Figure 6B; all values reported in Figure S3B). Among the UNF, POS, and NEG groups, all Primed cohorts consistently shared 6 over-expressed (>2-fold, *p*<0.05) and 6 under-expressed (<-2-fold, *p*<0.05) genes following an inflammatory challenge, building a basis of expected overall transcriptional results for the crude population. In descending order, these included increased gene expressions of *ICAM-1, IL-1β, IL-6, BMP-2, TNF-α,* and *LIF*, and in ascending order of down-regulated genes were *GDF5, COL1A1, BMP6, ITGAV, TGF-β3,* and *THY1*. These 12 genes were similarly altered suggesting that although these subpopulations are distinguishable in phenotype and function, both POS and NEG subpopulations fundamentally show similarities in most gene expressions in response to inflammatory challenge (Figures 6B-C). However, critical transcriptional differences were indeed identified that correlate with functions ascribed specifically to CD146^+^ BM-MSC. For instance, the master regulators and various other genes tightly related with osteogenic and adipogenic differentiation potential (*i.e., Runx2, PPARγ, BGLAP* and *ITGB1*), initially found downregulated in UNF Primed cells (Figure 3D), were found altered only in the NEG Primed group. This suggests that POS Primed cells, despite the inflammatory challenge, retain their molecular osteogenic and adipogenic machinery (Sacchetti et al., 2007; Yue et al., 2016) and that the suppression observed in the heterogeneous UNF Primed preparation can be attributed only to the NEG subpopulation. Furthermore, the POS Primed subpopulation showed additional signature gene expressions that revealed genes that may have larger implications. *BDNF* (3.58-fold; *p*<0.005) was over-expressed and *GDF15* (-4.06-fold; *p*<0.03) and *NUDT6* (-2.41-fold; *p*<0.01) were under-expressed in only the POS Primed subpopulation whereas no unique genes were related to the NEG subpopulations. Primed cohorts of the UNF and POS BM-MSC shared significantly up-regulated gene *VCAM1* and down-regulated genes *PDGF-Rβ* and *TGF-β1* following inflammatory challenge.

### Potency and immunomodulatory function of POS subpopulation support the signature profile with first responder characteristics

Potency associated with the POS or NEG subpopulations was determined, as with UNF cells, by the most robust suppression of proliferation of stimulated target cells using the IPA. Dosage responses using PBMC showed the most significant reduction in proliferation rates at 2:1 ratio of PBMC co-cultured with the POS Naïve (17.00±13.59%; *p*<0.001), POS Primed (7.75±8.54%; *p*<0.0001), NEG Naïve (16.25±14.61%; *p*<0.001), and NEG Primed (9.88±9.92%; *p*<0.001). More importantly, at 4:1 POS Naïve (39.00±17.38%; *p*<0.05) and POS Primed (5.50±5.20%; *p*<0.0001) significantly outperformed the immunosuppressive potential of NEG Naïve (36.13±30.93%) and NEG Primed (13.38±14.26%; *p*<0.001) subpopulations. Moreover, PBMC proliferation was significantly suppressed at 12:1 by the POS Primed (17.50±8.81%; *p*<0.001) and to lesser extent NEG Primed (22.63±20.37%; *p*<0.01) cohorts (Figure 7A).

**Figure 7.**
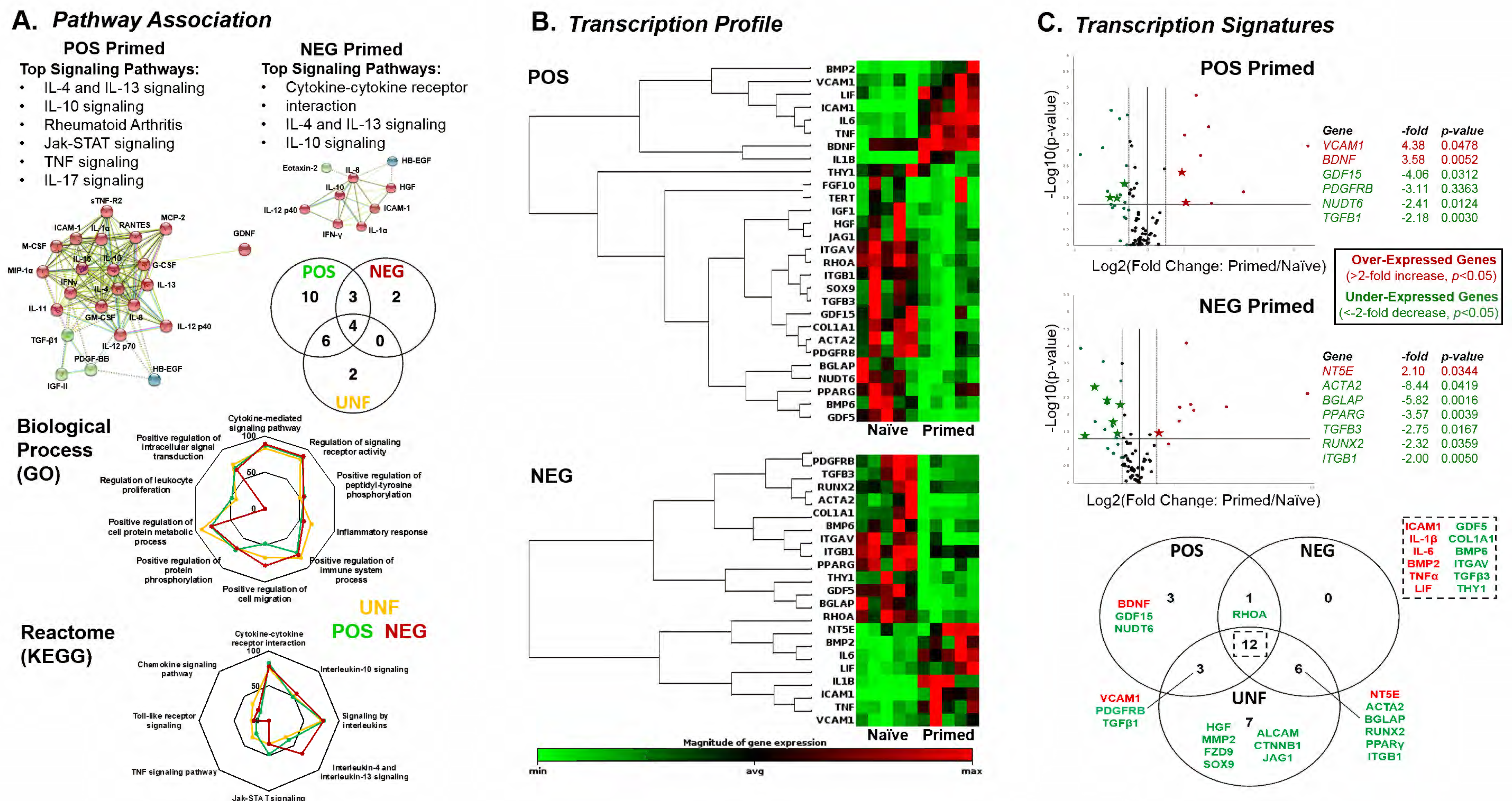
Immunopotency assays of POS and NEG cohorts reveal markedly robust potency of POS BM-MSC cohorts. A. Group comparison between UNF, POS and NEG groups showed a significant reduction of stimulated T cell proliferation by co-culture with POS Naïve BM-MSC than NEG Naïve. B. Proliferation rates of stimulated T cells co-cultured with Naïve and Primed cohorts of POS and NEG BM-MSC demonstrated the potency of the POS Naïve and POS Primed groups compared to the NEG Naïve group. C. Images of the 72-hour duration of the immunopotency assay showing the proliferation of the stimulated T cells (green) that are co-cultured above the adherent BM-MSC groups. D. T cell subsets differentially promoted by the co-culture with Naïve and Primed cohorts of the POS and NEG BM-MSC. E. The effects of the conditioned media (CM) from all groups modulated the differentiation of T cell subsets after 72 hours of culture.

Comparisons of the T cell IPA results from the Naïve cohorts of the UNF, POS, and NEG BM-groups demonstrated the intrinsic contributions of the subpopulations within the UNF BM-MSC, explaining the results presented in Figure 2B. The increased T cell proliferation observed with the UNF Naïve cohort (32.11±16.85%) can now be exclusively attributed to the NEG fraction. Unlike the T cell proliferation rate reduction to 16.72± 18.75% with the POS Naïve cohort (comparable to the 11.56±8.86% without cells, Figure 2B), a significantly higher rate was observed with the NEG Naïve cohort reaching 42.62±7.87% (*p*<0.01) (Figure 7B). Although donor-specific variations are observable, especially within the POS Naïve cohort, the reproducibility of this assay still showed a remarkable pattern among cohorts using the same donors in all groups. To further demonstrate the intrinsic immunosuppressive potency of the POS subpopulations, the POS Primed (12.81±15.91%) showed no difference to the POS Naïve cohort in T cell proliferation rates (Figures 7C-D and S4E), suggesting that the POS subpopulation is innately equipped to functionally immunomodulate independent of prior stimulation. The NEG Primed cohort was also able to reduce T cell proliferation (13.24±8.55%). However, when compared with the NEG Naïve rates, the necessity of a prior inflammatory challenge for this subset to functionally perform becomes evident.

As demonstrated with the UNF groups (Naïve and Primed), a detailed description of the subsets of differentiated T cells is necessary to obtain a more complete picture of the immunomodulatory effect of the POS and NEG subpopulations. POS Naïve (15.76±6.75%; *p*<0.05) and POS Primed (16.75±6.95%; *p*<0.01) BM-MSC groups coincided with markedly higher frequencies of Naïve T cells (CD3^+^CD4^-^CD8^-^CFSE^+^ cells) compared to controls (No BM-MSC; 7.16±3.14%), suggesting attenuation of T cell differentiation (Figure 7E and S4F). On the contrary, NEG Naïve (6.38±4.94%; *p*<0.01) and NEG Primed (5.55±4.84%; *p*<0.001) groups had no difference with the control but were significantly lower than their POS counterpart. The effects of all POS and NEG BM-MSC groups on Cytotoxic T cell differentiation were comparable to controls (30.35±17.03%), however a trend is observed with mild suppressive effects by the POS Naïve (21.51±11.59) and POS Primed (17.82±15.88%), not evident with the NEG Naïve (29.41±9.24) and NEG Primed (28.37±10.45%) cohorts. Similarly, POS Naïve (56.11±12.21%) and POS Primed (51.15±5.51%) BM-MSC modestly reduced differentiation of Helper T cells compared to controls (60.78±12.88%), while NEG Naïve (62.70±16.55%) and NEG Primed (64.14±15.74%) BM-MSC promoted modest differentiation into Helper T cell subsets. More importantly, analysis of the Treg frequency within the Helper T subset showed that POS Naïve (46.23±22.28%; *p*<0.0001) and POS Primed (66.66±28.06%; *p*<0.0001) induced the most significant amount compared to controls (5.56±3.87%). Further, induction by POS Naïve was markedly greater (*p*<0.01) than NEG Naïve BM-MSC (17.17±9.33%), and POS Primed was significantly higher (*p*<0.0001) than NEG Primed BM-MSC (16.30±6.50%; Figure 7E and S4F). Collectively, this compelling evidence underscores the mechanistic underpinnings of immunomodulation in which the POS subpopulation outperforms the NEG subpopulation in several approaches.

### POS subpopulation ameliorates synovitis and fat pad fibrosis by the induction of M2 alternative activation macrophages

In order to correlate the enhanced immunomodulatory properties found in the POS subpopulation with *in vivo* therapeutic efficacy, we used a well-established rat model of synovium and infrapatellar fat pad (IFP) inflammation and fibrosis (Takahashi et al., 2018a; Udo et al., 2016). In that model, the intra-articular administration of monosodium iodoacetate (MIA) rapidly induces local inflammatory changes leading to progressive thickening of synovium (*i.e.,* synovitis) and fibrosis throughout the IFP that can be quantified. Therefore, this model permits multiparameter measurements not only of synovium and IFP structural alterations but also of underlying local cellular changes driving them, all with promising clinical implications. As shown in the experimental outline (Figure 8A), synovitis/IFP fibrosis induction was followed 4 days later (at the peak of local inflammation (Takahashi et al., 2018b)) by a single intra-articular injection of BM-MSC (UNF, POS or NEG). Sham (no MIA and injection with vehicle only) and untreated (MIA induction followed by no treatment) animals were used as healthy and diseased controls, respectively. Evaluation of each group was performed by a standard semi-quantitative scoring method (Takahashi et al., 2018a; Udo et al., 2016), where synovitis and IFP fibrosis are independently assessed (0 to 3 for each). The latter is then complemented by a stain-based quantitative evaluation based on computational algorithm (Figure S4B).

**Figure 8.**
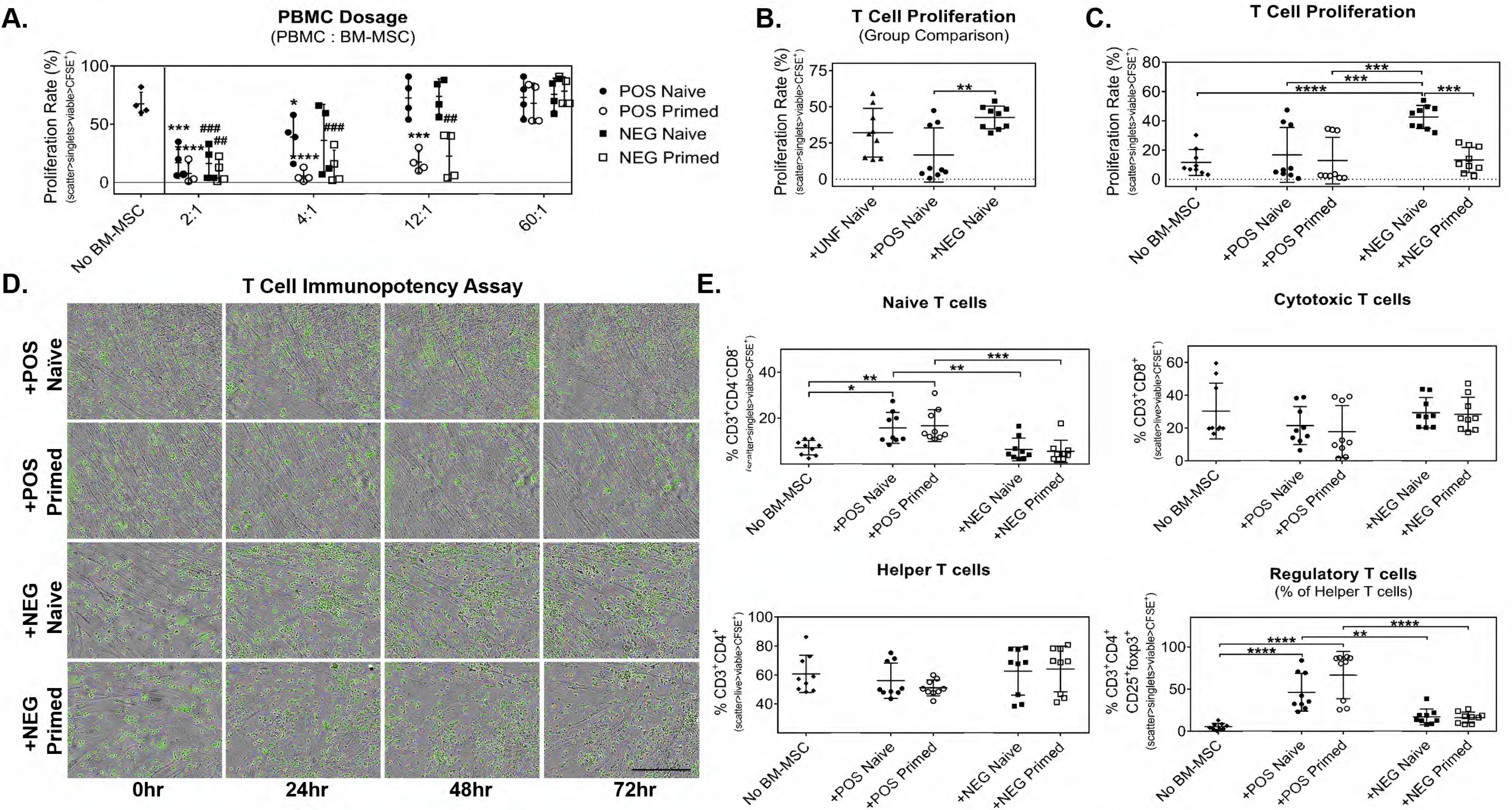
*In vivo* assessment of therapeutic efficacy of UNF, POS, and NEG BM-MSC following synovitis and IFP fibrosis of the knee. A. Experimental design of the *in vivo* induction of synovitis/IFP fibrosis, treatment with BM-MSC, and tissue assessments. B. H&E-stained sections of the synovium and IFP were imaged at 4X magnification for structural changes indicative of synovitis and IFP fibrosis. Asterisks and arrows denote areas of observable fibrosis and synovitis, respectively. C. Using a standard scoring system and computational approach, quantitative comparisons of synovitis and scoring were performed. Black data points signify female animals, blue data points signify male animals. Black asterisks denote statistical difference against Sham group, red asterisks denote differences against Untreated group. *, *p*<0.05; **, *p*<0.01; ***, *p*<0.001; ****, *p*<0.0001.

Synovitis scores showed a significant inflammatory reaction in treated animals (n=6) after the induction with MIA, evident when comparing Sham (0.17±0.24) with the Untreated (3.00±0.00; *p*<0.0001). Treatment with UNF (1.67±0.52; *p*<0.0001), POS (1.33±0.52; *p*<0.01), and NEG (2.25±0.52; *p*<0.0001) showed significantly greater synovitis scores compared to Sham with POS treatment with the greatest amelioration. This improvement was further demonstrated by the comparison of scores against the Untreated synovitis score with the POS (*p*<0.0001) and UNF (*p*<0.01) treatment markedly reduced while no difference was seen with the NEG treatment. Additionally, POS (*p*<0.05) had significantly lower synovitis score than the NEG treatment group. Similar trends were apparent with the IFP fibrosis scores showing significant differences in all groups (Untreated *p*<0.0001; UNF *p*<0.05; POS *p*<0.05; NEG *p*<0.001) compared to Sham (0.25±0.50) Compared to the Untreated group (2.75±0.50), treatment with UNF (1.5±0.63; *p*<0.05) and POS (1.5±0.45; *p*<0.05) BM-MSC were markedly reduced and no differences with NEG BM-MSC (2.25±0.76). Similar supporting data was obtained with the computational fibrosis quantification, which found a comparable trend as with the scoring system, with reduced fibrosis in UNF (0.33±0.05%), POS (0.29±0.14%) and less with NEG (0.39±0.21%) BM-MSC compared to the Untreated (0.45±0.10%). The clear trend in fibrosis evaluations, yet with no statistical differences, can be attributed to our early assessment time point (8 days after MIA induction). At this early point, unlike the established synovitis, it is expected that the fibrotic changes are variable. Furthermore, large differences were depicted between female and male recipients that contributed to the variability (Figure 8B-C).

To determine mechanistic cellular changes promoted locally by BM-MSC *in vivo*, M1/M2 macrophage phenotypic evaluations were performed (represented as the ratio of M2 over M1 for consistent comparisons), as MSC modulate macrophage polarization (reviewed in (Bernardo and Fibbe, 2013)). Synovium/IFP resident and infiltrated macrophages have demonstrated key roles in joint health and articular cartilage degenerative changes, as they are the source of inflammatory and catabolic mediators including IFNγ, TNFα, IL-1,-6,-8 and metalloproteinases (reviewed in (Bondeson et al., 2010)). Therefore, the modulation of the macrophage phenotype within those structures has significant consequences. Classically activated-M1 (CD86^+^) and alternatively activated-M2 (CD206^+^) macrophages were immunolocalized to the synovium and throughout the body of the IFP, specifically surrounding the vasculature (Figures 9A-B). Differences were observed between male and female recipients, thus the results presented separately by gender. Furthermore, in spite of variations among recipients, consistent trends were observed. In females, M2 and M1 macrophages were comparably observed within synovium and IFP tissues of the Sham group, resulting in a 0.98±0.95 M2/M1 ratio. Untreated recipients showed a reduced M2/M1 score of 0.53±0.11 demonstrating the balance tipped when largely pro-inflammatory M1 macrophages are present. More importantly, treatment with both UNF (1.68±0.98 M2/M1) and POS (1.13±1.00M2/M1) BM-MSC switched the ratios to positive values (*i.e.* M2 values larger than M1 values), suggesting active macrophage polarization towards an anti-inflammatory therapeutic M2 phenotype. This remarkable improvement was not observed with NEG BM-MSC treatment (0.54±0.47 M2/M1), which showed a comparable ratio to the Untreated group suggesting an inability to revert the M1 phenotype by this subpopulation (Figure 9A). Unlike females, the male Sham group (0.26±0.14 M2/M2) showed little presence of M2 macrophages compared to the high presence of M1, resulting in similar values to the Untreated IFP group (0.22±0.22 M2/M1). The remaining groups showed comparable results as with females, with a large induction of M2 macrophages with both UNF (1.53±2.06 M2/M1) and more so POS (1.86±2.24) BM-MSC treatment, and NEG BM-MSC (0.42±0.31 M2/M1) undistinguishable from Untreated (Figure 9B).

**Figure 9.**
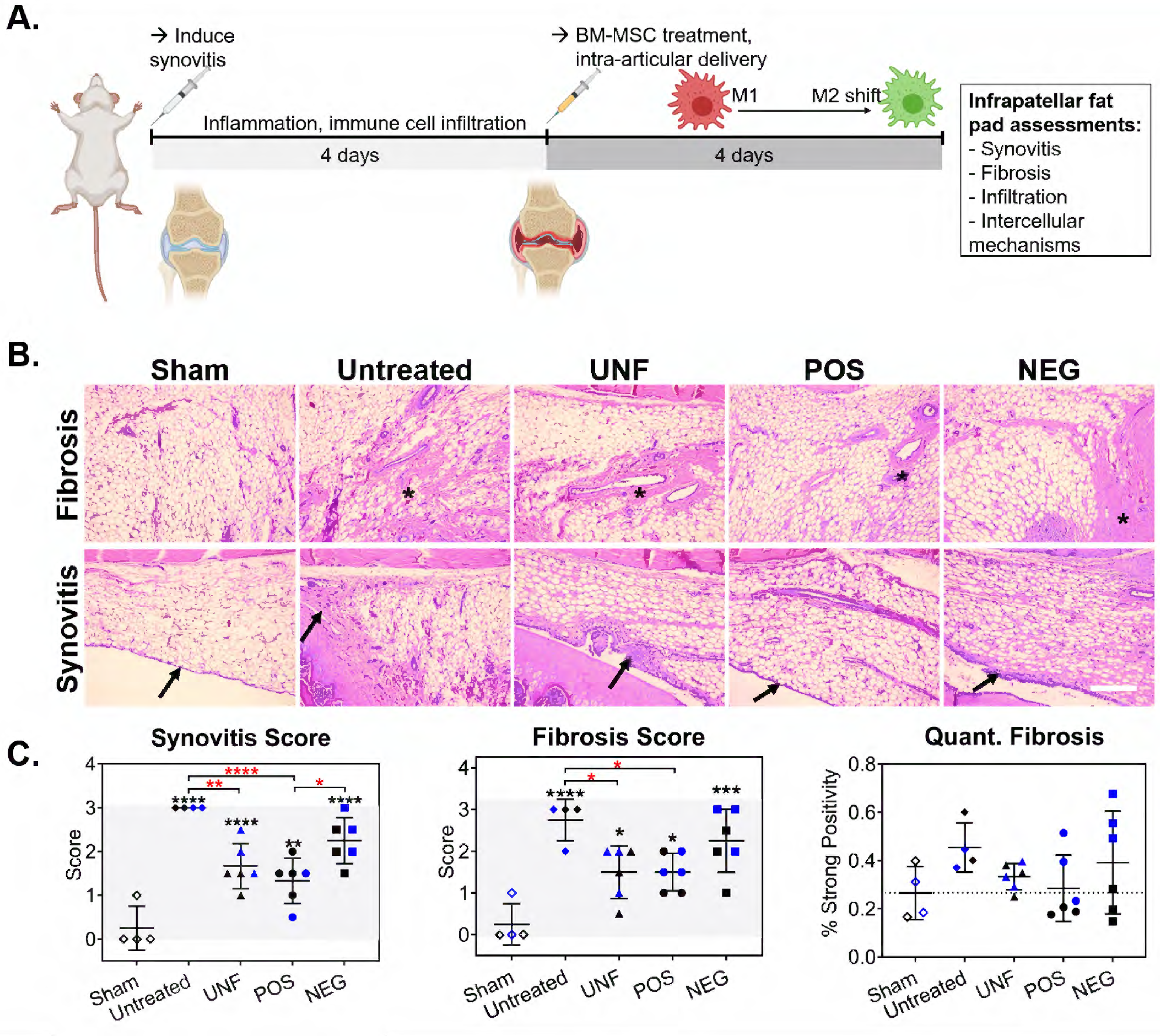
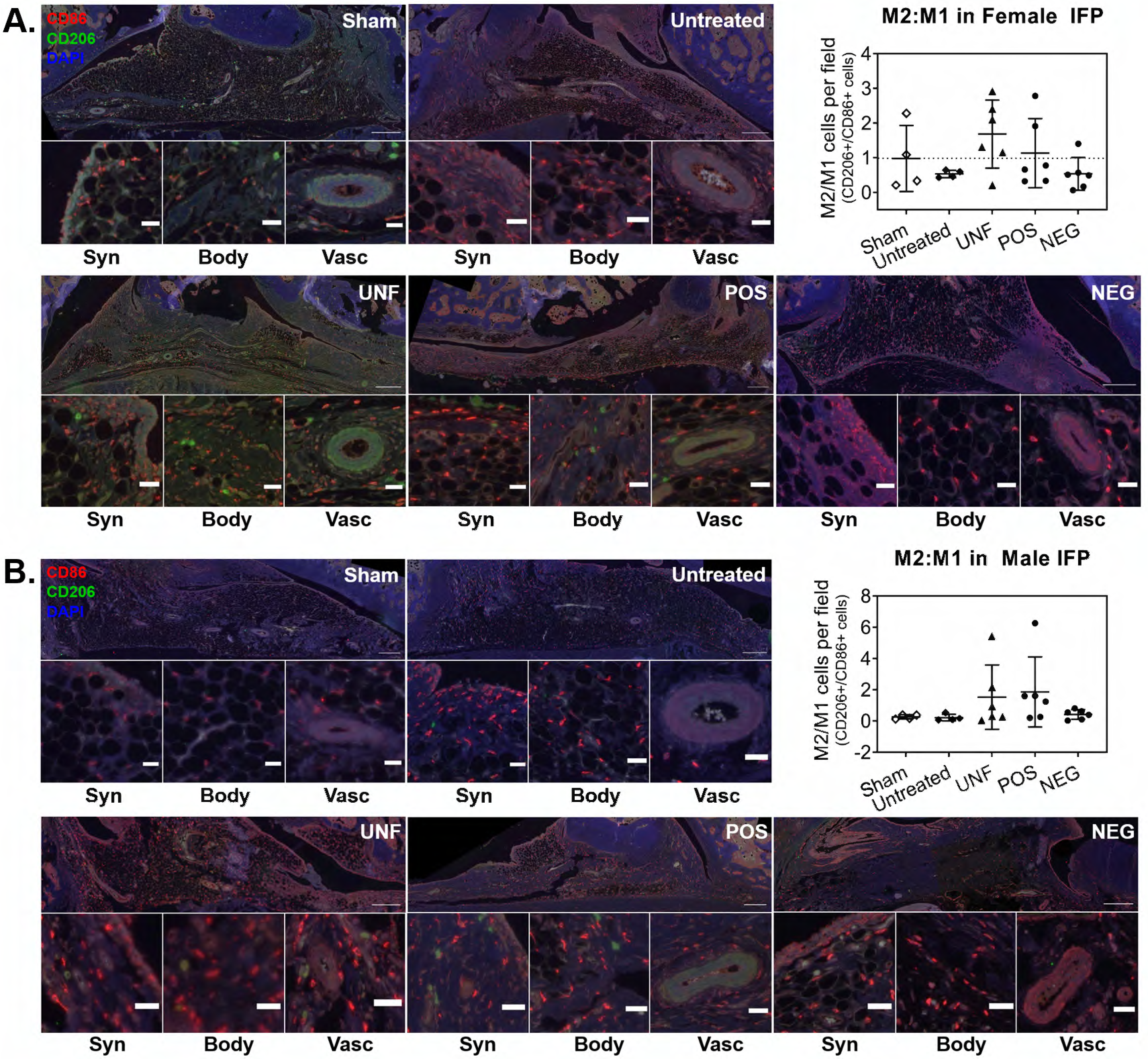
*In vivo* M1-to-M2 shift induction by POS in areas of synovitis and IFP fibrosis. Images and quantification for M1 (CD86; classical activation) and M2 (CD206; alternative activation) macrophages presence in female (A) and male (B) knee sections in each group. M2:M1 ratios define the positive effects following treatment representing a majority of M2 over M1 macrophages. Scale bars represent 500 µm (top images) and 100 µm (bottom images).

## DISCUSSION

The heterogeneity and identification of distinct subpopulations within crude MSC have been implicated to confound evidence and expected outcomes when used as therapeutic biologics (Buhring et al., 2009; Phinney, 2012; Siegel et al., 2013; Sivasubramaniyan et al., 2016; 2012). These effects slow advancements and engender regulatory concerns even when standard methodologies are practiced. Donor-specific differences (*i.e.,* age, sex, health status) also correlate to the “cell fitness” and therapeutic efficacy adding yet another complexity among donor comparisons (Frazier et al., 2018; Scruggs et al., 2013; Yu et al., 2011). Previous evidence has helped distinguishing subpopulations within crude MSC based on the expression of the pericyte-related marker CD146. These include descriptions of their developmental origins (Corselli et al., 2010; Maijenburg et al., 2012; Sacchetti et al., 2016; Slukvin and Kumar, 2018), association with the perivascular niche (Corselli et al., 2013b; 2013a; Covas et al., 2008; Crisan et al., 2008; Espagnolle et al., 2014; Tormin et al., 2011), self-renewal and stem-like capacities supporting hematopoiesis (Mendelson and Frenette, 2014b; Méndez-Ferrer et al., 2010; Sacchetti et al., 2016; 2007) and therapeutic use as osteoblastic progenitors for bone repair (James et al., 2017; 2012). Our results further support this emerging distinction, now beyond those progenitor and hematopoietic support contributions for the CD146^+^ (POS) subpopulation. Herein, we correlated in great detail phenotypic signatures and transcriptional/secretory differences in response to inflammatory challenge with enhanced functional potency based on immunomodulatory and anti-inflammatory effects.

Pericytes (*a.k.a.,* Rouget cells) have long been associated with structural functions in blood vessels including stabilization, tone control, angiogenesis and permeability (reviewed in (Gutiérrez et al., 2009)). Additional functions such as macrophage-like properties and effects on the immune system (*e.g.,* cell trafficking and responses) have now been more clearly elucidated (reviewed in (Navarro et al., 2016)). On the other hand, we have previously shown that perivascular BM-MSC function as gatekeepers during metastatic cancer cell extravasation, exercised through a CD146-mediated mechanism (Correa et al., 2016). Collectively, the strategic perivascular localization seems to furnish resident cells with sensing capabilities to survey, guide and instruct cells through trans-endothelial trafficking. Based on these observations, we hypothesized that CD146^+^ perivascular BM-MSC would more robustly influence immune/inflammatory cascades than their counterparts more distant from vascular structures (CD146^-^). Such a functional discrimination has immediate repercussions not only for our understanding of MSC biology, but also to streamline manufacturing protocols for cell-based products with less heterogeneous and more predictable compositions.

To experimentally test our hypothesis, we employed an *in vitro* technique of inflammatory challenge (*i.e.*, TNF-α, IFN-γ priming) to both interrogate phenotypic adaptations and examine the potency of the responses from each BM-MSC donor and associated subpopulations. The results obtained with UNF Naïve and Primed cohorts built the foundation for the subsequent evaluation of the CD146+ (POS) and CD146- (NEG) subpopulations, where correlative evidence suggested the contribution of each subpopulation within the whole UNF preparation. Cell priming has been used as a strategic method to incite an analyzable response by the cells or to pre-condition them to augment a desired trait or effect (Kadle et al., 2018; Silva et al., 2018). Moreover, priming BM-MSC prior to performing immunopotency assays using target cells better recapitulates the *in vivo* pathologic events involving active secretion of inflammatory molecules followed by cellular infiltration of immune cells (*e.g.,* T cells and whole PBMC) and activation of resident cells (*e.g.,* macrophages) typical of tissue damage (Julier et al., 2017). We found that priming UNF BM-MSC induced a consistent response among all 8 donors examined regardless of age and gender. This included a CD146^high^CD107a^high^CXCR4^high^LepR^high^ phenotype, reminiscent of the perisinusoidal MSC within the BM (Mendelson and Frenette, 2014a; Méndez-Ferrer et al., 2010) with highly sensing and secretory capacities. Remarkably, the sorted POS Primed subpopulation uniquely recapitulated this phenotypic enrichment (LepR increased but not significantly, though), suggesting that the overall phenotypic change seen in the UNF preparation can be attributed to the POS group.

The resulting phenotype of UNF Primed BM-MSC corresponded to a transcriptional profile that indicated a more stem-like and immunomodulatory state with increased key transcripts (*e.g., LIF*). Together with a highly immunomodulatory secretory profile (*e.g.,* PGE-2 and IDO activity), they were consistent with the results of the IPA that achieved robust suppression of PBMC proliferation (even at high PBMC:MSC ratios), overcoming the declining dose-dependent effect of UNF naïve cells. The deep analysis of the subpopulations’ secretory profiles revealed that the POS was most robustly in response to the inflammatory challenge, strongly suggesting that it is the POS subpopulation, and not the NEG, the potently responsive cell type contributing to the overall effect of the UNF BM-MSC population. This aligns with the enhanced secretory potency and the higher CD107a inducible expression found in the POS group. Functionally, the POS Primed contributed more than the NEG Primed cohort to the stabilization of the PBMC proliferation suppression even at high PBMC:MSC ratios, strengthening the concept of a robust immunosuppressive subpopulation.

The analysis of T cell proliferation and subset distribution showed additional compelling evidence for the immunomodulatory capacity of BM-MSC, and specifically the POS subpopulation. The stimulus used triggers the activation of T cell response, yet lacks perpetuating stimuli yielding only mild proliferation but induction of differentiation to the major subsets as main measurable outcomes. The expected additional proliferative response of T cells when non-stimulated UNF Naïve BM-MSC were present (W. Li et al., 2012) was explained by the effects of the NEG subpopulation exclusively, as the POS Naïve indeed suppressed T cell proliferation effectively. More interestingly, this intrinsic immunosuppressive influence of the Naïve cohort of the POS subpopulation eliminated the pre-exposure to pro-inflammatory molecules (*i.e.,* IFNγ-TNFα priming), previously reported as requisite to induce immunosuppressive activities of MSC via NO/IDO and monocyte presence (Bernardo and Fibbe, 2013; François et al., 2012; W. Li et al., 2012; Stagg and Galipeau, 2013). It also provides information to substantiate the phenotypic distinction of MSC into type 1 and 2, which largely depends on the absence or presence of an IFNγ/TNFα-rich environment they sense, respectively (Bernardo and Fibbe, 2013; Waterman et al., 2010). Priming demonstrated a marked suppression of T cell proliferation in all groups (*i.e.*, UNF Primed, POS Primed and NEG Primed) comparable to the effect of the POS Naïve group. Beyond these distinct effects on overall T cell proliferation, specific and significant differences were also generated by POS and NEG groups in the subset composition of T cells. The robust induction of undifferentiated naïve T cells and Tregs exclusive of the POS group, supports the fact that MSC are known to shift the T cell axis from cytotoxic to Tregs for reparative and tolerogenic effects (J. Li et al., 2018; Miyagawa et al., 2017). However, now clarifying the identity of the cells responsible for that shift (POS), and their innate potency to exert that effect even without prior stimulation/priming. An overall attenuation of T cell differentiation provides a fundamental mechanism of BM-MSC immunomodulatory effects necessary to translate for therapeutic use.

UNF Primed BM-MSC also showed downregulation of the multi-lineage differentiation master genes *RUNX2*, *SOX9*, and *PPARγ.* Priming, then, enabled a controlled shift toward stemness and immunomodulation and away from progenitor capabilities, which coincides with the inherent characteristic of asymmetrical division of MSC. However, the compromised differentiation programs (especially osteogenic and adipogenic) was initially difficult to reconcile with the observed CD146 and LepR enriched phenotype after priming. This, based on the osteogenic nature of CD146^+^ perivascular osteoprogenitors (Sacchetti et al., 2007) and perivascular stem/stromal cells (PSC) (James et al., 2017; 2012; Wang et al., 2019), as well as LepR^+^ adipogenic progenitors (Yue et al., 2016). The apparent dichotomy was elucidated after analyzing the transcriptional profiles of the sorted and primed POS and NEG subpopulations. Downregulation of *RUNX2* and *PPARγ*, along with other key transcripts responsible for healthy osteogenic and adipogenic programs were exclusively shared between UNF and NEG groups, leaving POS with no alterations in those molecular programs. Consequently, the selective reduction in differentiation potential of the NEG subpopulation suggests the POS as the functional subset capable of exerting either function depending on the environment they sense.

Relevant to translational efforts, all the *in vitro* functional discrimination between POS and NEG was successfully translated *in vivo*. Intra-articular injection of POS BM-MSC therapeutically outperformed NEG cohorts, greatly improving synovitis and IFP fibrosis within the joint, correlative to promoting rapid M2-macrophage phenotypic shift. Macrophages are receiving special attention given their orchestrating roles regulating local reparative changes, thus becoming therapeutic cell targets. In fact, a triangulation involving MSC, macrophages and T cells has been described by François *et al*. (François et al., 2012). They show that MSC-derived IDO activity drives the M2 polarization of macrophages, which, in turn help suppress T cell proliferation thus amplifying the MSC effect. On the other hand, M2 polarized macrophages counteract the secretion of inflammatory and catabolic molecules by the M1 type, reducing joint damage especially to the articular cartilage (Bartok and Firestein, 2010; Bondeson et al., 2010; Eymard et al., 2014). We have previously demonstrated the homing and transient engraftment (≤ 7 days) of MSC intra-articularly injected in this model of synovitis/IFP fibrosis (Kouroupis et al., 2019). Therefore, we propose that highly sensitive, secretory and potent immunosuppressive POS BM-MSC revert inflammatory and fibrotic changes in the synovium and IFP, in part via inducing a key reparative M2 macrophage phenotypic shift.

These high content data provided several methods of characterizing the functional potency of BM-MSC that can be further adapted for standardization in efforts to test potency across tissue-specific MSC preparations. Sorting methods to capture greater purity between subpopulations should be implemented to overcome the limitations of MACS. Although much of the data contains significant trends, deviations among donors stress the need to detect donor-specific differences to better tailor potential biologics as therapies. Further investigations into the functional potency may allow for greater understanding of functional subpopulations (*e.g.,* CD146^+^ BM-MSC) within the crude population of MSC, determining “cell fitness” representative to donor health, and ultimately predictivity of *in vivo* cell interactions for better alignment with expected outcomes.

### Conclusion

This study bolsters the notion of distinct MSC functional subpopulations based on CD146 expression, previously determined based on osteogenic potential and hematopoietic control and now elevated to immunomodulatory potential and therapeutic efficacy. Our data strongly supports the idea of the CD146^+^CD107a^+^ subpopulation harboring the robust sensory/secretory/immunomodulatory cells within crude preparations, deemed “first responders”. These findings have translational implications aimed to comply with rigor and transparency as well as indicates trends and tools adaptable for cell manufacturing by minimizing variability and offering a way to assess and predict the cell-based product’s potency.

## MATERIALS AND METHODS

### Human BM-MSC culture and priming with inflammation cocktail

Human BM-MSC were obtained from 8 consent-signed, de-identified patients and used for all experiments at passage 3 or 4 (demographic information; Figure S1A). Briefly, BM-MSC in complete culture media (CCM) containing of Gibco™ Dulbecco’s Modified Eagle Medium + Glutamax (Thermo Fisher Scientific, Inc., Waltham, MA) and 10% fetal bovine serum (Seradigm, VWR, Radnor, PA) were expanded as adherent cells and maintained in a humidified incubator set to 37°C and 5% CO2. Media was replaced every 3-4 days with fresh CCM until 70% confluence was achieved. BM-MSC were then washed with phosphate buffered saline (PBS) and incubated with TripLE EXPRESS (Thermo Fisher Scientific) for 5 minutes at 37°C. Upon detachment, BM-MSC were neutralized with CCM (v/v), collected, and pelleted by centrifugation (1500 rpm, 5 minutes). BM-MSC were counted using Trypan Blue (Thermo Fisher Scientific) dead cell exclusion method, and either directly assayed or stored at -80°C until further use.

BM-MSC used as crude cells were designated as the unfractionated (UNF) group. UNF BM-MSC were subsequently sorted based on CD146 expression to yield the CD146^+^ (POS) and CD146^-^ (NEG) subpopulations. Briefly, UNF BM-MSC were re-suspended in staining buffer containing PBS with 0.5% bovine serum albumin (BSA) and 2 mM EDTA and then incubated with biotinylated anti-human CD146 (Miltenyi Biotech, Inc., Auburn, CA) at 4°C for 20 minutes. Invitrogen™ CELLection Dynabeads™ Biotin Binder Kit (Thermo Fisher Scientific) were used according to manufacturer’s instructions for magnet-activated cell sorting resulting in the POS and NEG subpopulations. POS and NEG BM-MSC were directly placed in culture for a minimal culture period or directly plated for assays.

UNF, POS, or NEG BM-MSC were designated into cohorts, termed Naïve (*i.e.,* maintained in CCM) or Primed. Primed cohorts were exposed to an inflammation cocktail prepared with 10µg/ml IFNγ and 15µg/ml TNFα (R&D Systems, Minneapolis, MN) added to CCM. After 48 hours, Naïve and Primed BM-MSC and derived conditioned media (CM) were subsequently analyzed by the following assays.

### Phenotypic and transcriptional analyses

Phenotypic expression was assessed using a CytoFLEX flow cytometer (Beckman Coulter Life Sciences, Indianapolis, IN). Briefly, Naïve and Primed cohorts of UNF, POS, and NEG BM-MSC (n=8) were suspended in staining buffer and incubated with fluorescently conjugated anti-human antibodies detailed in Table S1 for 20 minutes at 4°C. Cells were then washed twice with staining buffer, fixed in 4% paraformaldehyde for 5 minutes at room temperature, following two additional washes. Cells were transferred to 4°C and acquisition of 10,000 events was performed within 3 days. Using Kaluza software (Beckman Coulter), samples were gated (scatter>singlets>viability) and positive gates were placed according to controls for desired phenotypic expressions. Overlays were generated to show expressions of representative samples with isotype and unstained controls.

Total RNA was isolated and purified from Naïve and Primed cohorts of UNF (Naïve and Primed, n=7), POS (Naïve and Primed, n=5), and NEG (Naïve and Primed, n=5) BM-MSC using RNeasy® Plus Mini Kits (Qiagen) according to the manufacturer’s instruction. RNA was measured, prepared as 1μg reactions using SuperScript™ VILO™ cDNA Synthesis Kit (Invitrogen), and transcribed to cDNA using a thermocycler. Following, real-time quantitative polymerase chain reactions (qPCR) were performed using RT2 Profiler Arrays for Human Mesenchymal Stem Cells (Qiagen). Briefly, SYBR Green Supermix, 100 µl cDNA, and Ultrapure water, and 20 µl mastermix was added into each well of the array plates. Ct values were obtained and oriented into Excel spreadsheets for importing into Qiagen’s Data Analysis Center where ΔCt was obtained using all included housekeeping genes, and ΔΔCt was calculated based on the Primed cohort normalized to the Naïve cohort of the same group. Data was represented as unsupervised hierarchical clustering of each group represented the relationships corresponding to gene expression activities, and relative fold changes were reported with statistical significance.

### Standard characterization of BM-MSC groups

BM-MSC were characterized for spindle-shaped morphology, growth kinetics, multi-lineage differentiation capacity, and phenotypic expression. For growth kinetics, Naïve and Primed cohorts of UNF BM-MSC (n=8) were seeded in triplicate into the wells of a 24-well plates and placed in an IncuCyte ZOOM system (Essen Bioscience, Inc., Ann Arbor, MI) set to 37°C and 5% CO_2_ set to 10X magnification for 8 days. Images were acquired for morphology and analysis of the proliferation rate using ZOOM software (Essen Bioscience, Inc.; Figure.) For induction of adipogenesis and osteogenesis, UNF Naïve BM-MSC (n=5) achieved 80-100% confluence and then incubated with adipogenic and osteogenic/chondrogenic differentiation media (StemPro™, Gibco), respectively, that was replaced every 3-4 days. After 21 days, cells were washed and stained with Oil Red O or Alizarin Red for the detection of lipid droplets or calcium deposition, respectively. Images were acquired at 4X for osteogenesis and 10X for adipogenesis using a Compact Leica CTR microscope with LAS X software (Leica Microsystems, Wetzler, Germany). Thereafter, elution of Oil Red O or Alizarin Red with isopropyl or 10% cetylpyridinium chloride solution, respectively, was measured using a plate reader at OD584. Remaining de-stained samples were lysed and measured for total protein using Pierce™ BCA Protein Assay Kit (Thermo Fisher Scientific) according to the manufacturer’s instructions. Quantification of adipogenesis and osteogenesis was calculated by the detected OD of stain divided by the total protein of each sample, represented as the normalized OD. Chondro-pellet cultures, 0.25x10^6^ UNF Naïve BM-MSC (n=3) were induced towards chondrogenesis for 21 days with serum-free MesenCult-ACF differentiation medium (STEMCELL Technologies Inc, Vancouver, Canada). Harvested pellets were cryosectioned and 6-μm frozen sections stained with 1% toluidine blue (Sigma).

### Immunocytochemistry for signature marker expression

Immunocytochemistry was performed to detect the expression of CD146, CD107a, CXCR4, and LepR on Naïve and Primed cohorts of UNF, POS, and NEG BM-MSC using antibodies detailed in Table S1. Briefly, BM-MSC were washed with PBS and fixed in 10% formalin for 10 minutes at room temperature followed by 3 washes with PBS. Cells were subsequently incubated in in blocking solution containing 10% normal goat serum and 1% BSA in PBS for 30 minutes. Primary antibodies (1:200) were added to staining buffer containing 10% blocking solution in 1X Tris buffered saline (Bio-Rad Laboratories, Inc., Hercules, CA), and added to fixed cell samples for 1 hour at room temperature. Slides were washed with TBS and then incubated with PE-conjugated secondary antibodies in staining buffer for 1 hour. Following, cell samples were washed with TBS, mounted with Prolong Gold with DAPI (Thermo fisher Scientific), and imaged at 10X magnification.

### Secretome analysis by enzyme-linked immunosorbent assays (ELISA)

CM was collected from Naïve or Primed cohorts of UNF, POS, and NEG BM-MSCs from 48 hours of culture in CCM (naïve cohorts) or inflammation cocktail (primed cohort). CM samples (n=7) were analyzed for the presence of PGE2 using Prostaglandin E2 ELISA Kit (Cayman Chemical, Ann Arbor, MI) according to the manufacturer’s instructions. Additionally, Human C-Series ELISA Inflammation and Growth Factor Arrays (RayBiotech Life, Inc., Norcross, GA) were performed according to the manufacturer’s instructions to detect soluble molecules in the CM of each sample (n=8 for all samples). Using the provided excel-based analysis plug-in, all samples were normalized by background subtraction and quantitatively compared by normalization of the Primed cohort with the corresponding Naïve cohort. Protein interactomes were generated by Search Tool for Retrieval of Interacting Genes/Proteins (STRING 11.0; http://string-db.org) database using interaction data from experiments, databases, neighbourhood in genome, gene fusions, co-occurrence across genomes, co-expression and text-mining. An interaction confidence score of 0.4 was imposed to ensure high interaction probability whereas K-means clustering algorithm organized proteins into 3 separate clusters per condition tested, discriminated by colors. Venn diagrams were used to demonstrate all possible relations between UNF, POS, and NEG BM-MSCs for the significantly (*p<0.05*) altered proteins. Functional enrichments related to biological process, Kyoto Encyclopedia of Genes and Genomes (KEGG) pathways, and reactome pathways were presented in radar graphs for all BM-MSC primed cohorts.

### Indoleamine 2,3-Dioxygenase (IDO) Activity

Naïve and Primed cohorts of the UNF (n=6), POS (n=5), and NEG (n=5) groups were collected as pellets and used to measure the activity of IDO. According to the manufacturer’s protocol, Indoleamine 2,3-Dioxygenase 1 Activity Assay Kit (Abcam) was performed using cell lysates of each sample. Using a standard curve for *N*-formylkynurenine, quantitative analysis of metabolized tryptophan (1 mole:1 mole) as the byproduct of IDO activity was measured and normalized to the total amount of protein for each sample. Total protein was measured using Pierce™ BCA Protein Assay Kit (Thermo Fisher Scientific) provided by the standards and colorimetric detection using a plate reader.

### Analysis of secretory potency

Molecular Probes® CellTrace™ CFSE Cell Proliferation Kit (Thermo Fisher Scientific) was prepared as a working solution in PBS, and POS and NEG BM-MSC groups (n=4) were labeled by incubation at 37°C for 10 minutes in the dark. Stain was quenched by adding cold CCM, and BM-MSC were centrifuged, washed, and plated in CCM at a seeding density of 100,000 cells/well of 12-well plates. As controls, unlabeled BM-MSC were plated simultaneously for each donor. After 24 hours, fresh CCM was replaced in all conditions, and inflammation cocktail was added to designated Primed cohorts. Media from each condition was sampled in duplicate and read at excitation/emission wavelengths of 485/521nm using a plate reader. The sampling volume was replaced with fresh CCM with or without inflammation cocktail for Naïve or Primed cohorts, respectively, and sampling was performed at 1, 2, 4, 8, 12, 24, and 48 hours. Absorbance readings from unlabeled BM-MSC supernatants were used as background and subtracted from all sample values. All values for each donor POS or NEG BM-MSC were normalized to the corresponding Naïve cohort (represented as 1) for each time point. Additionally, values were represented as the area under curve comparison for the total 48 hours.

### Immunopotency assays (IPA)

UNF, POS, or NEG BM-MSC (n=3) were seeded as technical triplicates using 24-well plates. After 24-48 hours, media was replaced with either fresh CCM or inflammation cocktail (10µg/ml IFNγ and 15µg/ml TNFα) for BM-MSC cohorts designated as Naïve or Primed, respectively. Concomitantly, human Pan T cells (STEMCELL Technologies) were thawed or peripheral blood mononuclear cells (PBMCs; IRB#19950119) were isolated and subsequently cultured in flasks with complete RPMI media containing RPMI supplemented with 15% Human Serum AB (Corning), 1% 1mM Sodium Pyruvate, 1% 0.1mM Non-essential Amino Acid, 1% 1X Vitamins, 1% 10nM HEPES, and 1% 2mM L-glutamine (Thermo Fisher Scientific). After 48 hours of priming, BM-MSC were washed and media was replaced with complete RPMI media. Simultaneously, T cells or PBMCs were stained with CellTrace™ CFSE Cell Proliferation Kit as previously described. Following, cells were counted using live/dead exclusion method, and T cells or PBMCs were directly co-cultured with BM-MSC groups (2:1). Dosage responses were demonstrated with co-cultures prepared with PBMCs to BM-MSC groups at ratios of 2:1, 4:1, 12:1, or 60:1. Following, 25µL of ImmunoCult (STEMCELL Technologies) or PMA/Ionomycin (eBioscience, Thermo Fisher Scientific) was added to the wells designated for T cell or PBMC stimulation, respectively, and co-cultures were maintained for 72 hours. All cells were cultured in a humidified incubator set to 37°C and 5% CO_2_.

### IPA flow cytometric staining and analysis

For IPA containing T cells, phenotypic analysis of T cells and BM-MSC groups (2:1) was performed after the 72 hours of co-cultures. T cells were collected together with BM-MSCs for each condition. Non-adherent T cells were carefully collected into 15 mL conical tubes, corresponding wells were washed with PBS which was also collected in the tubes containing T cells, and wells with remaining adherent BM-MSC groups were then incubated with TryPLE EXPRESS for 5 minutes at 37°C. Complete RPMI was then added to the lifted BM-MSC, and BM-MSC were collected and added to tubes with corresponding T cells. Together, cells were centrifuged and re-suspended in staining buffer and incubated with antibody cocktails (Table S1) for 20 minutes at 4°C. Cells were washed with staining buffer and either incubated with 1% paraformaldehyde for 5 minutes at room temperature or fixation solution (Thermo Fisher Scientific) for 20 minutes at 4°C. Cells fixed in paraformaldehyde were washed twice with staining buffer, re-suspended in a final volume of 200 µl, and transferred to a 96-well plate. Cells in fixation solution were washed twice with 1X permeabilization buffer (Thermo Fisher Scientific) and incubated with antibodies against intracellular markers for 40 minutes at 4°C. Following, cells were washed twice with permeabilization buffer, re-suspended in staining buffer to a final volume of 200 µl, and transferred to a 96-well plate. Using a Cytoflex LS (Beckman Coulter, Inc., Brea, CA) and CytExpert software, at least 20,000 events were acquired from each sample. Analysis was performed using Kaluza software version 3.0. BM-MSCs were gated based on scatter, singlets, and then discriminated using live/dead gating. T cells were gated based on scatter, singlets, live/dead discrimination, and then positivity for CFSE. T cell proliferation rates (%) were calculated as: ((CFSE^LOW^ events/ CFSE^+^) x100). Remaining data was represented as the percent gated (%) of the reported phenotype.

For IPA containing PBMCs, PBMCs only were collected and analyzed by flow cytometry. PBMCs co-cultured at 2:1 with BM-MSC groups were collected and res-suspended in staining buffer and subsequently stained with antibody cocktails (Table S1) for 20 minutes at 4°C. Cells were then washed twice, re-suspended to a final volume of 200 µl, and transferred to a 96-well plate for analysis. PBMCs co-cultured for dose-responses were collected, stained with Ghost Red Viability dye only, and analyzed. Using a Cytoflex LS (Beckman Coulter, Inc., Brea, CA) and CytExpert software, 50,000 events were acquired from each sample. Analysis was performed using Kaluza software version 3.0. PBMCs were gated based on scatter, singlets, live/dead discrimination, and then positivity for CFSE. PBMC proliferation rates were calculated as: ((CFSE^LOW^ events/ CFSE^+^) x100).

### In Vivo Evaluation Using Acute Synovitis/IFP fibrosis Rat Model

All animal procedures were approved by the Institutional Animal Care and Use Committee at the University of Miami (approval #16-008-ad03) and were in compliance with the ARRIVE guidelines [citation]. 10-week-old female (n=3) and male (n=3) Sprague Dawley rats were housed one animal per cage and acclimated for 1 week after arrival with *ad libitum* food and water. Animals were housed in sanitary, ventilated rooms with controlled temperature, humidity, and 12-hour light/dark cycles.

Animals were anesthetized by isoflurane inhalation, and acute synovial/infrapatellar fat pad (IFP) inflammation/fibrosis was induced by intra-articular (i.a.) injection of 1 mg of mono-iodoacetate (MIA) in 50 µl saline. Knees were flexed at 90°, and MIA was injected into the medial side of the joint with a 27 G needle using the patellar ligament and articular line as anatomical references. Short exposure to MIA has shown to induce inflammation to the synovium and adjacent IFP (Takahashi et al., 2018a; Udo et al., 2016). After 3 days, a single dose of 500,000 UNF, POS, or NEG BM-MSC in Euro-Collins solution (Media Tech) was administered via i.a. injection using the same injection technique. As controls, Sham animals were induced with i.a. injection of Euro-Collins solution only, *i.e.,* no MIA, and Untreated animals were administered i.a. injection of MIA followed by administration of Euro-Collins solution only (no BM-MSC).

After 4 days, all animals were sacrificed, and knees were harvested and stored in 10% formalin. Knees were subsequently cleaned of tissue and placed in fresh formalin for complete fixation. Following, knees were processed by decalcification, paraffin-embedded, and sectioned using a microtome into 5-micron sections and mounted onto glass slides. Histologic analysis was performed after de-paraffinization and re-hydration. Sections were stained with hematoxylin and eosin (H&E) or fluorescently conjugated antibodies described in Table S1. For immunohistochemical staining, histological sections were incubated with universal blocking solution for 30 minutes at room temperature. Following, sections were incubated with primary antibodies CD86 and CD206 (1:500) overnight at 4° C. Sections were then washed with TBS and incubated with fluorescently conjugated secondary antibodies for 2 hours at room temperature. After several washes with TBS, sections were mounted with Prolong Gold with DAPI and coverslipped. For chemiluminescence, sections were incubated with anti-human mitochondria primary antibody for 4 hours at room temperature, washed with TBS, and then incubated with antibodies and solutions provided in HRP/DAB (ABC) detection IHC kit (Abcam, Cambridge, MA) according to the manufacturer’s instructions. Mayer’s hematoxylin was used as a counter stain and cells were mounted and then coverslipped. All cells were imaged at 10X or 20X magnification for further analysis.

### Histological analysis

H&E-stained sections were imported into Aperio ImageScope software (Leica Biosystems) and analyzed using an algorithm based on color intensity defined by the following input parameters: Hue Value 0.1, Hue Width 0.99, ColorSaturation Threshold 0.16, Iwp(High) 220, Iwp(Low) = Ip (High) 180, Ip(Low) = Isp(High) 100, and Inp(High) -1. Total histologic analysis was performed in 2 sections per animal (2 female controls, 2 male controls, 3 female treated, or 3 male treated), assessed for percent positive pixel quantification for fibrosis, and pixel-based measurements were obtained synovial membrane thickness using ImageScope software tools. Immunofluorescence images were imported into Fiji/ImageJ software (NIH, Bethesda, MA), and analysis was performed on 2 sections per animal. Cells captured in each channel (FITC, PE, or DAPI) were counted after tuning for cell only capture and represented as M2/M1 measured by cells count for M2 divided by M1.

### Statistical Analysis

All values for Naïve and Primed cohorts of UNF, POS, and NEG BM-MSC were reported as the average ± standard deviation with points plotted by each donor. Statistical analyses were performed using one-way or two-way analysis of variance (ANOVA) followed by pairwise comparisons using Tukey *post hoc* testing. Significance for the overall group effect and individual pairwise comparisons were made amongst the groups or against the Naïve cohort for each Primed cohort as described in the figure legends. All analysis was performed using Prism v7 (Graphpad Software, La Jolla, CA).

## ACKNOWLEDMENTS

We give great thanks to Arnold I. Caplan, Ph.D. and Rodrigo Somoza, Ph.D. for donating healthy donor BM-MSC used in our studies, and Oliver Umland, Ph.D. and Marcia Boulina, PhD for their technical assistance in the Diabetes Research Institute flow cytometry and imaging core facilities. We also thank Thomas M. Best, M.D., Ph.D for reviewing the manuscript.

## SUPPLEMENTARY FIGURES

**Figure S1.**
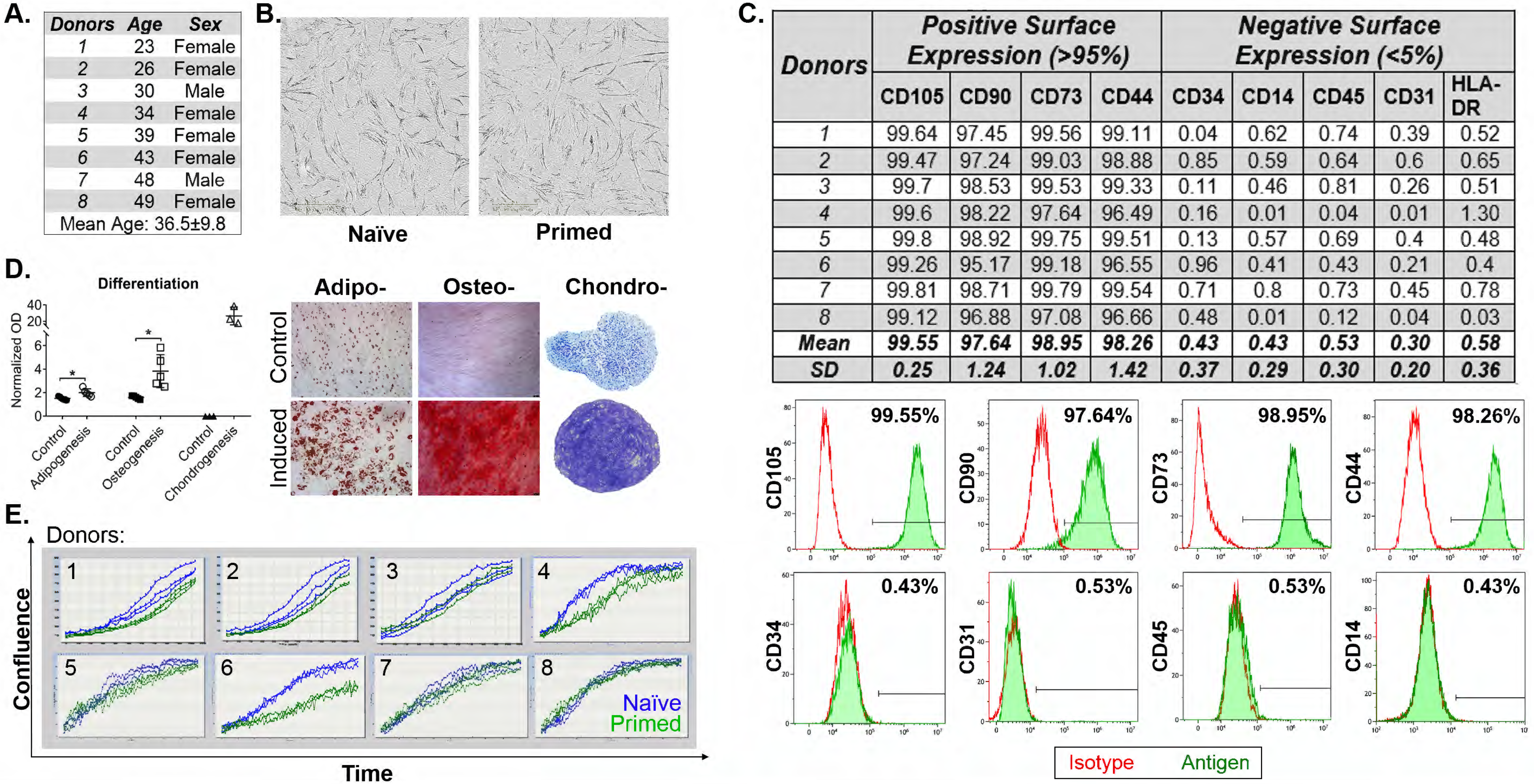
Characterization of human BM-MSC. A. Demographic information of each donor used which included both sexes of various ages. B-C. Multilineage differentiation capacity of BM-MSC induced with adipogenic, osteogenic and chondrogenic medias. D. Image of spindle-shape morphology indicative of BM-MSC. Scale bar represents 250 µm. E. Proliferation rate of BM-MSC to determine population doublings. F. Phenotypic analysis of positive and negative markers of each donor.

**Figure S2.**
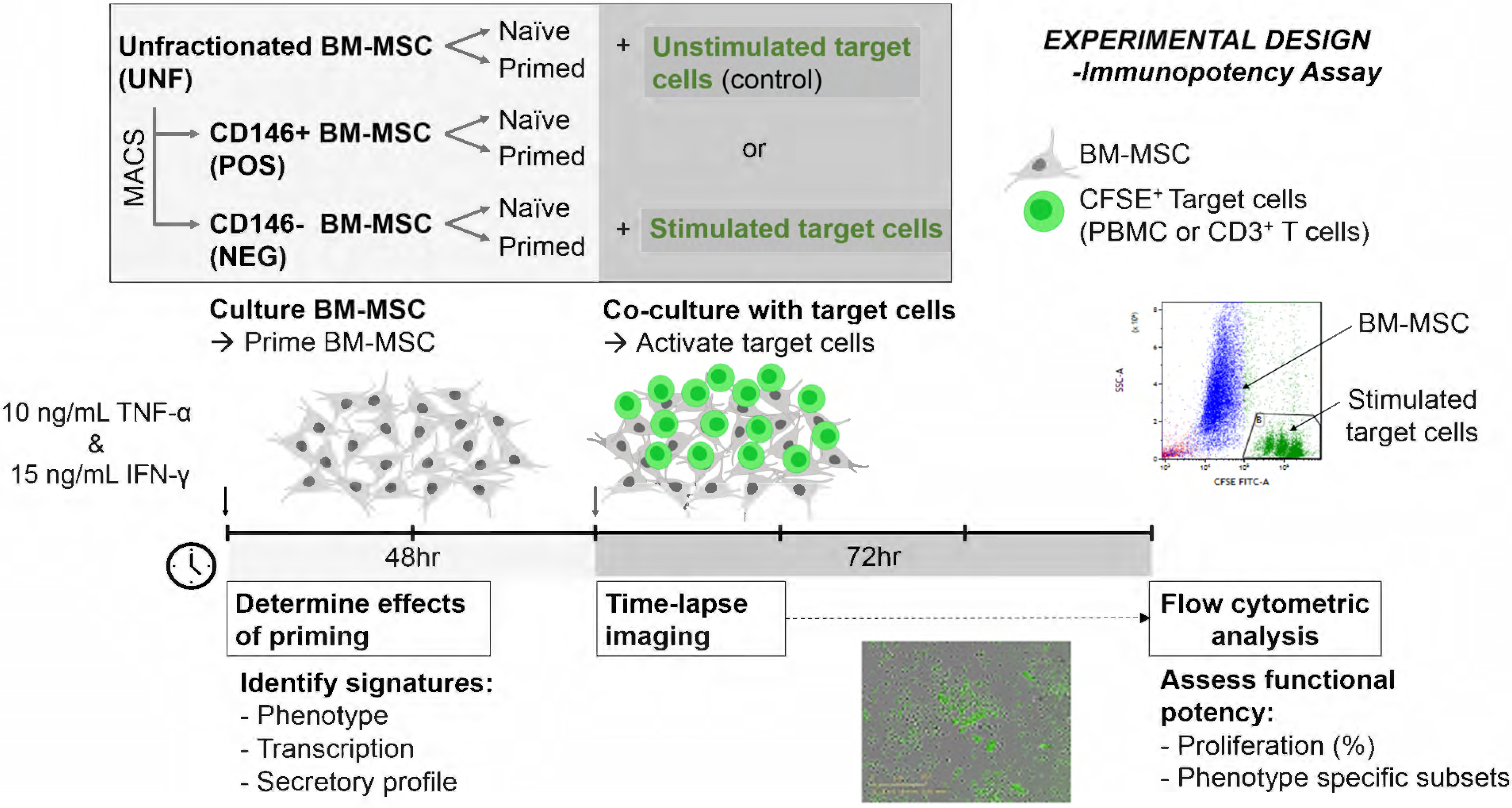
Experimental design for the immunopotency assay. Co-cultures were performed with Naïve or Primed cohorts of UNF, POS, or NEG BM-MSC and target cells, either stimulated T cells or PBMCs according to the timeline.

**Figure S3.**
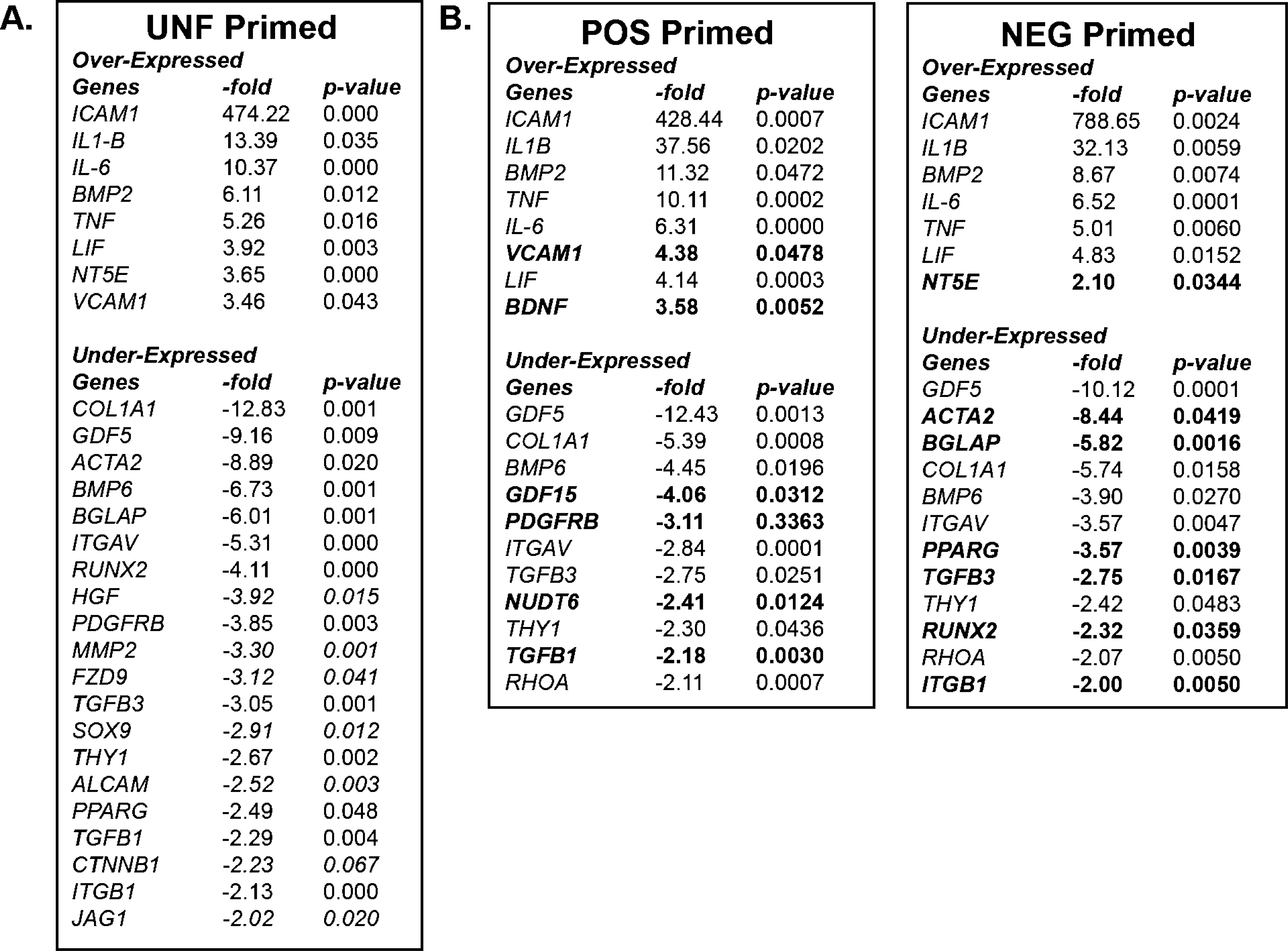
Inflammatory challenge deduced significantly altered gene transcripts. A. Gene expression analysis of UNF Primed cohorts identified over- (>2-fold) and under-expressed (<1-2-fold) transcripts of interest that were significantly altered upon inflammatory challenge. B. POS Primed or NEG Primed cohorts showed similar transcripts as UNF Primed that were significantly over- and under-expressed upon inflammatory challenge as well as distinguished transcripts specific to each subpopulation (bold).

**Figure S4.**
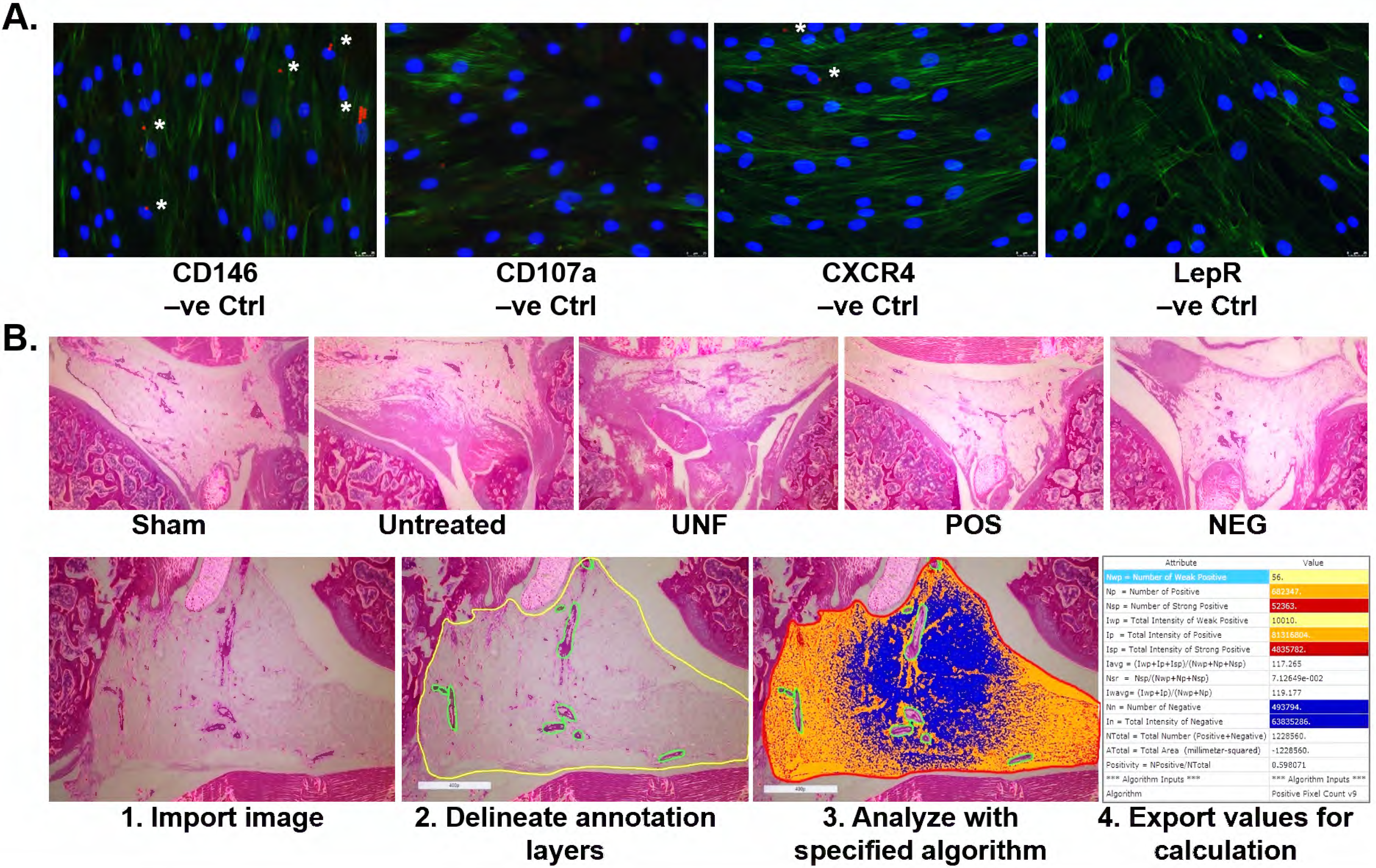
Analytic controls and parameters. A. Immunofluorescence negative controls for signature surface antigens CD146, CD107a, CXCR4, and LepR with DAPI (blue) and F-actin (green) stain. White areas show MACS beads engulfed by sorted cells that are fluorescent in the PE (red) channel while imaging. B. Histologic analysis of each group using H&E-stained sections. Quantitative processing of images was performed by: 1. Importing images into ImageScope software, 2. Delineating area of interest by creating annotation layers of included (yellow) and excluded (green) areas, 3. Running specified algorithm to analyze pixels positive for infiltration of cells and dense areas of fibrosis, and 4. Exporting values for calculation of the positive pixel count (%) of the total tissue area.

**Table S1:** Antibody information.

**4. Export values for calculation**
| Antibody | Clone | Manufacturer | Application |
| --- | --- | --- | --- |
| Ghost Dye™ Red780 Viability Dye |  | Tonbo Bioscience | FC |
| Isotype IgG2b | RTK4530 | Biolegend | FC |
| Isotype IgG2a | S43.10 | Miltenyi Biotec | FC |
| Isotype IgG1 | MOPC-21 | Biolegend | FC |
| Isotype IgG2a | eBM2a | BD Biosciences | FC |
| Isotype IgG1 | X40 | BD Biosciences | FC |
| Isotype IgG1 | MOPC-21 | BD Biosciences | FC |
| Isotype IgG2a | MOPC-173 | BD Biosciences | FC |
| Isotype IgG2a | G155-178 | BD Biosciences | FC |
| CD73 | AD2 | Biolegend | FC |
| CD90 | 5E10 | Biolegend | FC |
| CD44 | IM7 | Biolegend | FC |
| CD105 | SN6h | Biolegend | FC |
| CD45 | HI30 | BD Biosciences | FC |
| CD31 | WM59 | BD Biosciences | FC |
| CD34 | AC136 | Miltenyi Biotec | FC |
| CD14 | 61D3 | Thermo Fisher Scientific | FC |
| CD3e* | UCHT1 | BD Biosciences | FC |
| CD4 | OKT4(OKT4) | Thermo Fisher Scientific | FC |
| CD8 | RPA-T8 | Thermo Fisher Scientific | FC |
| foxp3* | PCH101 | Thermo Fisher Scientific | FC |
| CD25 | M-A251 | BD Biosciences | FC |
| CD86 | IT2.2 | Thermo Fisher Scientific | FC |
| CD206 | 19.2 | Thermo Fisher Scientific | FC |
| CD146 | 541-10B2 | Miltenyi Biotec | FC |
| CD146 | P1H12 | BD Biosciences | FC |
| CD107a (LAMP1) | H4A3 | Biolegend | FC |
| CD184 (CXCR4) | 12G5 | Invitrogen | FC |
| NG2 | 9.2.27 | BD Biosciences | FC |
| CD295 (Leptin R) | REA361 | Miltenyi Biotec | FC |
| HLA-DR | G46-6 | BD Biosciences | FC |
| CD146 | EPR3207(2) | Abcam | ICC |
| CXCR4 | UMB2 | Abcam | ICC |
| LAMP1 | Polyclonal | Abcam | ICC |
| NG2 | Polyclonal | Abcam | ICC |
| CD86 | BU63 | <u>Abcam</u> | ICC |
| CD206 | Polyclonal | <u>Abcam</u> | ICC |

\*intracellular staining method was performed for these markers

## REFERENCES

Alter, G., Malenfant, J.M., Altfeld, M., 2004. CD107a as a functional marker for the identification of natural killer cell activity. J. Immunol. Methods 294, 15–22. doi:10.1016/j.jim.2004.08.008

Bartok, B., Firestein, G.S., 2010. Fibroblast-like synoviocytes: key effector cells in rheumatoid arthritis. Immunol. Rev. 233, 233–255. doi:10.1111/j.0105-2896.2009.00859.x

Bernardo, M.E., Fibbe, W.E., 2013. Mesenchymal stromal cells: sensors and switchers of inflammation. 13, 392–402. doi:10.1016/j.stem.2013.09.006

Bondeson, J., Blom, A.B., Wainwright, S., Hughes, C., Caterson, B., van den Berg, W.B., 2010. The role of synovial macrophages and macrophage-produced mediators in driving inflammatory and destructive responses in osteoarthritis. Arthritis Rheum 62, 647–657. doi:10.1002/art.27290

Buhring, H.-J., Treml, S., Cerabona, F., de Zwart, P., Kanz, L., Sobiesiak, M., 2009. Phenotypic characterization of distinct human bone marrow-derived MSC subsets. Ann N Y Acad Sci 1176, 124–134. doi:10.1111/j.1749-6632.2009.04564.x

Caplan, A.I., 2017a. New MSC: MSCs as pericytes are Sentinels and gatekeepers. J Orthop Res 35, 1151–1159. doi:10.1002/jor.23560

Caplan, A.I., 2017b. Mesenchymal Stem Cells: Time to Change the Name! Stem Cells Transl Med 6, 1445–1451. doi:10.1002/sctm.17-0051

Caplan, A.I., Correa, D., 2011. The MSC: an injury drugstore. 9, 11–15. doi:10.1016/j.stem.2011.06.008

Caplan, A.I., Sorrell, J.M., 2015. The MSC curtain that stops the immune system. Immunol. Lett. 168, 136–139. doi:10.1016/j.imlet.2015.06.005

Chan, C.K.F., Gulati, G.S., Sinha, R., Tompkins, J.V., Lopez, M., Carter, A.C., Ransom, R.C., Reinisch, A., Wearda, T., Murphy, M., Brewer, R.E., Koepke, L.S., Marecic, O., Manjunath, A., Seo, E.Y., Leavitt, T., Lu, W.-J., Nguyen, A., Conley, S.D., Salhotra, A., Ambrosi, T.H., Borrelli, M.R., Siebel, T., Chan, K., Schallmoser, K., Seita, J., Sahoo, D., Goodnough, H., Bishop, J., Gardner, M., Majeti, R., Wan, D.C., Goodman, S., Weissman, I.L., Chang, H.Y., Longaker, M.T., 2018. Identification of the Human Skeletal Stem Cell. Cell 175, 43–56.e21. doi:10.1016/j.cell.2018.07.029

Chinnadurai, R., Rajan, D., Qayed, M., Arafat, D., Garcia, M., Liu, Y., Kugathasan, S., Anderson, L.J., Gibson, G., Galipeau, J., 2018. Potency Analysis of Mesenchymal Stromal Cells Using a Combinatorial Assay Matrix Approach. CellReports 22, 2504–2517. doi:10.1016/j.celrep.2018.02.013

Correa, D., Somoza, R.A., Lin, P., Schiemann, W.P., Caplan, A.I., 2016. Mesenchymal stem cells regulate melanoma cancer cells extravasation to bone and liver at their perivascular niche. Int J Cancer 138, 417–427. doi:10.1002/ijc.29709

Corselli, M., Chen, C.-W., Crisan, M., Lazzari, L., Péault, B., 2010. Perivascular ancestors of adult multipotent stem cells. Arterioscler Thromb Vasc Biol 30, 1104–1109. doi:10.1161/ATVBAHA.109.191643

Corselli, M., Chen, C.-W., Sun, B., Yap, S., Rubin, J.P., Péault, B., 2012. The tunica adventitia of human arteries and veins as a source of mesenchymal stem cells. Stem Cells Dev 21, 1299–1308. doi:10.1089/scd.2011.0200

Corselli, M., Chin, C.J., Parekh, C., Sahaghian, A., Wang, W., Ge, S., Evseenko, D., Wang, X., Montelatici, E., Lazzari, L., Crooks, G.M., Peault, B., 2013a. Perivascular support of human hematopoietic stem/progenitor cells. Blood 121, 2891–2901. doi:10.1182/blood-2012-08-451864

Corselli, M., Crisan, M., Murray, I.R., West, C.C., Scholes, J., Codrea, F., Khan, N., Péault, B., 2013b. Identification of perivascular mesenchymal stromal/stem cells by flow cytometry. Cytometry A 83, 714–720. doi:10.1002/cyto.a.22313

Covas, D.T., Panepucci, R.A., Fontes, A.M., Silva, W.A., Orellana, M.D., Freitas, M.C.C., Neder, L., Santos, A.R.D., Peres, L.C., Jamur, M.C., Zago, M.A., 2008. Multipotent mesenchymal stromal cells obtained from diverse human tissues share functional properties and gene-expression profile with CD146+ perivascular cells and fibroblasts. Exp Hematol 36, 642–654. doi:10.1016/j.exphem.2007.12.015

Crisan, M., Yap, S., Casteilla, L., Chen, C.-W., Corselli, M., Park, T.S., Andriolo, G., Sun, B., Zheng, B., Zhang, L., Norotte, C., Teng, P.-N., Traas, J., Schugar, R., Deasy, B.M., Badylak, S., Buhring, H.-J., Giacobino, J.-P., Lazzari, L., Huard, J., Péault, B., 2008. A perivascular origin for mesenchymal stem cells in multiple human organs. 3, 301–313. doi:10.1016/j.stem.2008.07.003

da Silva Meirelles, L., Caplan, A.I., Nardi, N.B., 2008. In Search of the In Vivo Identity of Mesenchymal Stem Cells. STEM CELLS 26, 2287–2299. doi:10.1634/stemcells.2007-1122

da Silva Meirelles, L., Sand, T.T., Harman, R.J., Lennon, D.P., Caplan, A.I., 2009. MSC frequency correlates with blood vessel density in equine adipose tissue. Tissue Eng Part A 15, 221–229. doi:10.1089/ten.tea.2008.0103

Dominici, M., Le Blanc, K., Mueller, I., Slaper-Cortenbach, I., Marini, F., Krause, D., Deans, R., Keating, A., Prockop, D., Horwitz, E., 2006. Minimal criteria for defining multipotent mesenchymal stromal cells. The International Society for Cellular Therapy position statement. Cytotherapy 8, 315–317. doi:10.1080/14653240600855905

Ducy, P., Amling, M., Takeda, S., Priemel, M., Schilling, A.F., Beil, F.T., Shen, J., Vinson, C., Rueger, J.M., Karsenty, G., 2000. Leptin inhibits bone formation through a hypothalamic relay: a central control of bone mass. Cell 100, 197–207. doi:10.1016/s0092-8674(00)81558-5

Espagnolle, N., Guilloton, F., Deschaseaux, F., Gadelorge, M., Sensebé, L., Bourin, P., 2014. CD146 expression on mesenchymal stem cells is associated with their vascular smooth muscle commitment. J Cell Mol Med 18, 104–114. doi:10.1111/jcmm.12168

Eymard, F., Pigenet, A., Citadelle, D., Flouzat-Lachaniette, C.H., Poignard, A., Benelli, C., Berenbaum, F., Chevalier, X., Houard, X., 2014. Induction of an inflammatory and prodegradative phenotype in autologous fibroblast-like synoviocytes by the infrapatellar fat pad from patients with knee osteoarthritis. Arthritis Rheumatol 66, 2165–2174. doi:10.1002/art.38657

François, M., Romieu-Mourez, R., Li, M., Galipeau, J., 2012. Human MSC suppression correlates with cytokine induction of indoleamine 2,3-dioxygenase and bystander M2 macrophage differentiation. Mol Ther 20, 187–195. doi:10.1038/mt.2011.189

Frazier, T., Lee, S., Bowles, A., Semon, J., Bunnell, B., Wu, X., Gimble, J., 2018. Gender and age-related cell compositional differences in C57BL/6 murine adipose tissue stromal vascular fraction. Adipocyte 7, 183–189. doi:10.1080/21623945.2018.1460009

Friedman, J.M., Halaas, J.L., 1998. Leptin and the regulation of body weight in mammals. Nature 395, 763–770. doi:10.1038/27376

Gao, W., Thompson, L., Zhou, Q., Putheti, P., Fahmy, T.M., Strom, T.B., Metcalfe, S.M., 2009. Treg versus Th17 lymphocyte lineages are cross-regulated by LIF versus IL-6. Cell Cycle 8, 1444–1450. doi:10.4161/cc.8.9.8348

Gomes, J.P., Coatti, G.C., Valadares, M.C., Assoni, A.F., Pelatti, M.V., Secco, M., Zatz, M., 2018. Human Adipose-Derived CD146+ Stem Cells Increase Life Span of a Muscular Dystrophy Mouse Model More Efficiently than Mesenchymal Stromal Cells. DNA Cell Biol. 37, 798–804. doi:10.1089/dna.2018.4158

Gutiérrez, R., Madrid, J.F., Varela, H., Valladares, F., Acosta, E., Martín-Vasallo, P., Díaz-Flores, L., 2009. Pericytes. Morphofunction, interactions and pathology in a quiescent and activated mesenchymal cell niche. Histol. Histopathol. 24, 909–969. doi:10.14670/HH-24.909

Horwitz, E.M., Le Blanc, K., Dominici, M., Mueller, I., Slaper-Cortenbach, I., Marini, F.C., Deans, R.J., Krause, D.S., Keating, A., International Society for Cellular Therapy, 2005. Clarification of the nomenclature for MSC: The International Society for Cellular Therapy position statement. Cytotherapy. doi:10.1080/14653240500319234

James, A.W., Hindle, P., Murray, I.R., West, C.C., Tawonsawatruk, T., Shen, J., Asatrian, G., Zhang, X., Nguyen, V., Simpson, A.H., Ting, K., Péault, B., Soo, C., 2017. Pericytes for the treatment of orthopedic conditions. Pharmacol. Ther. 171, 93–103. doi:10.1016/j.pharmthera.2016.08.003

James, A.W., Zara, J.N., Zhang, X., Askarinam, A., Goyal, R., Chiang, M., Yuan, W., Chang, L., Corselli, M., Shen, J., Pang, S., Stoker, D., Wu, B., Ting, K., Péault, B., Soo, C., 2012. Perivascular stem cells: a prospectively purified mesenchymal stem cell population for bone tissue engineering. Stem Cells Transl Med 1, 510–519. doi:10.5966/sctm.2012-0002

Jiao, J., Milwid, J.M., Yarmush, M.L., Parekkadan, B., 2011. A mesenchymal stem cell potency assay. Methods Mol Biol 677, 221–231. doi:10.1007/978-1-60761-869-0_16

Julier, Z., Park, A.J., Briquez, P.S., Martino, M.M., 2017. Promoting tissue regeneration by modulating the immune system. Acta Biomater 53, 13–28. doi:10.1016/j.actbio.2017.01.056

Kadle, R.L., Abdou, S.A., Villarreal-Ponce, A.P., Soares, M.A., Sultan, D.L., David, J.A., Massie, J., Rifkin, W.J., Rabbani, P., Ceradini, D.J., 2018. Microenvironmental cues enhance mesenchymal stem cell-mediated immunomodulation and regulatory T-cell expansion. PLoS ONE 13, e0193178. doi:10.1371/journal.pone.0193178

Kannan, K., Stewart, R.M., Bounds, W., Carlsson, S.R., Fukuda, M., Betzing, K.W., Holcombe, R.F., 1996. Lysosome-associated membrane proteins h-LAMP1 (CD107a) and h-LAMP2 (CD107b) are activation-dependent cell surface glycoproteins in human peripheral blood mononuclear cells which mediate cell adhesion to vascular endothelium. Cell. Immunol. 171, 10–19. doi:10.1006/cimm.1996.0167

Kehl, D., Generali, M., Mallone, A., Heller, M., Uldry, A.-C., Cheng, P., Gantenbein, B., Hoerstrup, S.P., Weber, B., 2019. Proteomic analysis of human mesenchymal stromal cell secretomes: a systematic comparison of the angiogenic potential. NPJ Regen Med 4, 8–13. doi:10.1038/s41536-019-0070-y

Kouroupis, D., Bowles, A.C., Willman, M.A., Perucca Orfei, C., Colombini, A., Best, T.M., Kaplan, L.D., Correa, D., 2019. Infrapatellar fat pad-derived MSC response to inflammation and fibrosis induces an immunomodulatory phenotype involving CD10-mediated Substance P degradation. Sci Rep 9, 10864. doi:10.1038/s41598-019-47391-2

Kouroupis, D., Sanjurjo-Rodriguez, C., Jones, E., Correa, D., 2018a. Mesenchymal Stem Cell Functionalization for Enhanced Therapeutic Applications. Tissue Eng Part B Rev ten.teb.2018.0118. doi:10.1089/ten.teb.2018.0118

Kouroupis, D., Sanjurjo-Rodriguez, C., Jones, E., Correa, D., 2018b. MSC functionalization for enhanced therapeutic applications. Tissue Eng Part B Rev ten.TEB.2018.0118. doi:10.1089/ten.TEB.2018.0118

Krampera, M., Galipeau, J., Shi, Y., Tarte, K., Sensebé, L., MSC Committee of the International Society for Cellular Therapy (ISCT), 2013. Immunological characterization of multipotent mesenchymal stromal cells--The International Society for Cellular Therapy (ISCT) working proposal. Cytotherapy 15, 1054–1061. doi:10.1016/j.jcyt.2013.02.010

Laschober, G.T., Brunauer, R., Jamnig, A., Fehrer, C., Greiderer, B., Lepperdinger, G., 2009. Leptin receptor/CD295 is upregulated on primary human mesenchymal stem cells of advancing biological age and distinctly marks the subpopulation of dying cells. Exp. Gerontol. 44, 57–62. doi:10.1016/j.exger.2008.05.013

Li, J., Tan, J., Martino, M.M., Lui, K.O., 2018. Regulatory T-Cells: Potential Regulator of Tissue Repair and Regeneration. Front Immunol 9, 585. doi:10.3389/fimmu.2018.00585

Li, M., Yu, J., Li, Y., Li, D., Yan, D., Ruan, Q., 2010. CXCR4+ progenitors derived from bone mesenchymal stem cells differentiate into endothelial cells capable of vascular repair after arterial injury. Cellular reprogramming 12, 405–415. doi:10.1089/cell.2009.0088

Li, W., Ren, G., Huang, Y., Su, J., Han, Y., Li, J., Chen, X., Cao, K., Chen, Q., Shou, P., Zhang, L., Yuan, Z.-R., Roberts, A.I., Shi, S., Le, A.D., Shi, Y., 2012. Mesenchymal stem cells: a double-edged sword in regulating immune responses. Cell Death Differ 19, 1505–1513. doi:10.1038/cdd.2012.26

Liu, X., Duan, B., Cheng, Z., Jia, X., Mao, L., Fu, H., Che, Y., Ou, L., Liu, L., Kong, D., 2011. SDF-1/CXCR4 axis modulates bone marrow mesenchymal stem cell apoptosis, migration and cytokine secretion. Protein & cell 2, 845–854. doi:10.1007/s13238-011-1097-z

Maijenburg, M.W., Kleijer, M., Vermeul, K., Mul, E.P.J., van Alphen, F.P.J., van der Schoot, C.E., Voermans, C., 2012. The composition of the mesenchymal stromal cell compartment in human bone marrow changes during development and aging. Haematologica 97, 179–183. doi:10.3324/haematol.2011.047753

Mendelson, A., Frenette, P.S., 2014a. Hematopoietic stem cell niche maintenance during homeostasis and regeneration. Nat Med 20, 833–845.

Mendelson, A., Frenette, P.S., 2014b. Hematopoietic stem cell niche maintenance during homeostasis and regeneration. Nat Med 20, 833–846. doi:10.1038/nm.3647

Metcalfe, S.M., 2011. LIF in the regulation of T-cell fate and as a potential therapeutic. Genes Immun. 12, 157–168. doi:10.1038/gene.2011.9

Metcalfe, S.M., Strom, T.B., Williams, A., Fahmy, T.M., 2015. Multiple Sclerosis and the LIF/IL-6 Axis: Use of Nanotechnology to Harness the Tolerogenic and Reparative Properties of LIF. Nanobiomedicine (Rij) 2, 5. doi:10.5772/60622

Méndez-Ferrer, S., Michurina, T.V., Ferraro, F., Mazloom, A.R., Macarthur, B.D., Lira, S.A., Scadden, D.T., Ma’ayan, A., Frenette, P.S., 2010. Mesenchymal and haematopoietic stem cells form a unique bone marrow niche. Nature 466, 829–834. doi:10.1038/nature09262

Miyagawa, I., Nakayamada, S., Nakano, K., Yamagata, K., Sakata, K., Yamaoka, K., Tanaka, Y., 2017. Induction of Regulatory T Cells and Its Regulation with Insulin-like Growth Factor/Insulin-like Growth Factor Binding Protein-4 by Human Mesenchymal Stem Cells. J Immunol 199, 1616–1625. doi:10.4049/jimmunol.1600230

Nasef, A., Mazurier, C., Bouchet, S., François, S., Chapel, A., Thierry, D., Gorin, N.C., Fouillard, L., 2008. Leukemia inhibitory factor: Role in human mesenchymal stem cells mediated immunosuppression. Cell. Immunol. 253, 16–22. doi:10.1016/j.cellimm.2008.06.002

Navarro, R., Compte, M., Álvarez-Vallina, L., Sanz, L., 2016. Immune Regulation by Pericytes: Modulating Innate and Adaptive Immunity. Front Immunol 7, 480. doi:10.3389/fimmu.2016.00480

Niu, C.C., Lin, S.S., Chen, W.J., Liu, S.J., Chen, L.H., Yang, C.Y., Wang, C.J., Yuan, L.J., Chen, P.H., Cheng, H.Y., 2015. Benefits of biphasic calcium phosphate hybrid scaffold-driven osteogenic differentiation of mesenchymal stem cells through upregulated leptin receptor expression. Journal of orthopaedic surgery and research 10, 111. doi:10.1186/s13018-015-0236-2

Phinney, D.G., 2012. Functional heterogeneity of mesenchymal stem cells: implications for cell therapy. J Cell Biochem 113, 2806–2812. doi:10.1002/jcb.24166

Sacchetti, B., Funari, A., Michienzi, S., Di Cesare, S., Piersanti, S., Saggio, I., Tagliafico, E., Ferrari, S., Robey, P.G., Riminucci, M., Bianco, P., 2007. Self-renewing osteoprogenitors in bone marrow sinusoids can organize a hematopoietic microenvironment. Cell 131, 324–336. doi:10.1016/j.cell.2007.08.025

Sacchetti, B., Funari, A., Remoli, C., Giannicola, G., Kogler, G., Liedtke, S., Cossu, G., Serafini, M., Sampaolesi, M., Tagliafico, E., Tenedini, E., Saggio, I., Robey, P.G., Riminucci, M., Bianco, P., 2016. No Identical “Mesenchymal Stem Cells“” at Different Times and Sites: Human Committed Progenitors of Distinct Origin and Differentiation Potential Are Incorporated as Adventitial Cells in Microvessels. Stem Cell Reports 6, 897–913. doi:10.1016/j.stemcr.2016.05.011

Salgado, A.J., Gimble, J.M., 2013. Secretome of mesenchymal stem/stromal cells in regenerative medicine. Biochimie 95, 2195. doi:10.1016/j.biochi.2013.10.013

Salgado, A.J., Gimble, J.M., Costa, B.M., 2018. The cell secretome in personalized and regenerative medicine. Biochimie 155, 1. doi:10.1016/j.biochi.2018.11.004

Salgado, A.J., Sousa, J.C., Costa, B.M., Pires, A.O., Mateus-Pinheiro, A., Teixeira, F.G., Pinto, L., Sousa, N., 2015. Mesenchymal stem cells secretome as a modulator of the neurogenic niche: basic insights and therapeutic opportunities. Front Cell Neurosci 9, 249. doi:10.3389/fncel.2015.00249

Scheller, E.L., Song, J., Dishowitz, M.I., Soki, F.N., Hankenson, K.D., Krebsbach, P.H., 2010. Leptin functions peripherally to regulate differentiation of mesenchymal progenitor cells. STEM CELLS 28, 1071–1080. doi:10.1002/stem.432

Schwab, K.E., Gargett, C.E., 2007. Co-expression of two perivascular cell markers isolates mesenchymal stem-like cells from human endometrium. Hum. Reprod. 22, 2903–2911. doi:10.1093/humrep/dem265

Scruggs, B.A., Semon, J.A., Zhang, X., Zhang, S., Bowles, A.C., Pandey, A.C., Imhof, K.M.P., Kalueff, A.V., Gimble, J.M., Bunnell, B.A., 2013. Age of the donor reduces the ability of human adipose-derived stem cells to alleviate symptoms in the experimental autoimmune encephalomyelitis mouse model. Stem Cells Transl Med 2, 797–807. doi:10.5966/sctm.2013-0026

Siegel, G., Kluba, T., Hermanutz-Klein, U., Bieback, K., Northoff, H., Schäfer, R., 2013. Phenotype, donor age and gender affect function of human bone marrow-derived mesenchymal stromal cells. BMC Med 11, 146. doi:10.1186/1741-7015-11-146

Silva, L.H.A., Antunes, M.A., Santos, Dos, C.C., Weiss, D.J., Cruz, F.F., Rocco, P.R.M., 2018. Strategies to improve the therapeutic effects of mesenchymal stromal cells in respiratory diseases. Stem Cell Res Ther 9, 45. doi:10.1186/s13287-018-0802-8

Sivasubramaniyan, K., Harichandan, A., Boss, P., Buehring, H.J., van Osch, G., 2016. Isolation of phenotypically and functionally distinct endogeneous human bone marrow-derived mesenchymal stem/stromal cell subsets. Osteoarthr Cartil 24, S464. doi:10.1016/j.joca.2016.01.846

Sivasubramaniyan, K., Lehnen, D., Ghazanfari, R., Sobiesiak, M., Harichandan, A., Mortha, E., Petkova, N., Grimm, S., Cerabona, F., de Zwart, P., Abele, H., Aicher, W.K., Faul, C., Kanz, L., Buhring, H.-J., 2012. Phenotypic and functional heterogeneity of human bone marrow– and amnion-derived MSC subsets. Ann N Y Acad Sci 1266, 94–106. doi:10.1111/j.1749-6632.2012.06551.x

Slukvin, I.I., Kumar, A., 2018. The mesenchymoangioblast, mesodermal precursor for mesenchymal and endothelial cells. Cell Mol Life Sci. doi:10.1007/s00018-018-2871-3

Solchaga, L.A., Zale, E.A., 2012. Prostaglandin E2: a putative potency indicator of the immunosuppressive activity of human mesenchymal stem cells. American journal of stem cells 1, 138–145.

Stagg, J., Galipeau, J., 2013. Mechanisms of immune modulation by mesenchymal stromal cells and clinical translation. Curr. Mol. Med. 13, 856–867.

Sudworth, A., Dai, K.Z., Vaage, J.T., Kveberg, L., 2016. Degranulation Response in Cytotoxic Rat Lymphocytes Measured with a Novel CD107a Antibody. Front Immunol 7, 572. doi:10.3389/fimmu.2016.00572

Takahashi, I., Matsuzaki, T., Kuroki, H., Hoso, M., 2018a. Induction of osteoarthritis by injecting monosodium iodoacetate into the patellofemoral joint of an experimental rat model. PLoS ONE 13, e0196625–15. doi:10.1371/journal.pone.0196625

Takahashi, I., Matsuzaki, T., Kuroki, H., Hoso, M., 2018b. Induction of osteoarthritis by injecting monosodium iodoacetate into the patellofemoral joint of an experimental rat model. PLoS ONE 13, e0196625. doi:10.1371/journal.pone.0196625

Tomchuck, S.L., Zwezdaryk, K.J., Coffelt, S.B., Waterman, R.S., Danka, E.S., Scandurro, A.B., 2008. Toll-Like Receptors on Human Mesenchymal Stem Cells Drive Their Migration and Immunomodulating Responses. STEM CELLS 26, 99–107. doi:10.1634/stemcells.2007-0563

Tormin, A., Li, O., Brune, J.C., Walsh, S., Schütz, B., Ehinger, M., Ditzel, N., Kassem, M., Scheding, S., 2011. CD146 expression on primary nonhematopoietic bone marrow stem cells is correlated with in situ localization. Blood 117, 5067–5077. doi:10.1182/blood-2010-08-304287

Trickett, A., Kwan, Y.L., 2003. T cell stimulation and expansion using anti-CD3/CD28 beads. J. Immunol. Methods 275, 251–255.

Udo, M., Muneta, T., Tsuji, K., Ozeki, N., Nakagawa, Y., Ohara, T., Saito, R., Yanagisawa, K., Koga, H., Sekiya, I., 2016. Monoiodoacetic acid induces arthritis and synovitis in rats in a dose- and time-dependent manner: proposed model-specific scoring systems. Osteoarthr Cartil 24, 1284–1291. doi:10.1016/j.joca.2016.02.005

Vego, H., Sand, K.L., Hoglund, R.A., Fallang, L.E., Gundersen, G., Holmoy, T., Maghazachi, A.A., 2016. Monomethyl fumarate augments NK cell lysis of tumor cells through degranulation and the upregulation of NKp46 and CD107a. Cellular & molecular immunology 13, 57–64. doi:10.1038/cmi.2014.114

Wang, Y., Xu, J., Chang, L., Meyers, C.A., Zhang, L., Broderick, K., Lee, M., Péault, B., James, A.W., 2019. Relative contributions of adipose-resident CD146+ pericytes and CD34+ adventitial progenitor cells in bone tissue engineering. NPJ Regen Med 4, 1. doi:10.1038/s41536-018-0063-2

Waterman, R.S., Morgenweck, J., Nossaman, B.D., Scandurro, A.E., Scandurro, S.A., Betancourt, A.M., 2012. Anti-inflammatory mesenchymal stem cells (MSC2) attenuate symptoms of painful diabetic peripheral neuropathy. Stem Cells Transl Med 1, 557–565. doi:10.5966/sctm.2012-0025

Waterman, R.S., Tomchuck, S.L., Henkle, S.L., Betancourt, A.M., 2010. A new mesenchymal stem cell (MSC) paradigm: polarization into a pro-inflammatory MSC1 or an Immunosuppressive MSC2 phenotype. PLoS ONE 5, e10088. doi:10.1371/journal.pone.0010088

Wattrang, E., Dalgaard, T.S., Norup, L.R., Kjærup, R.B., Lundén, A., Juul-Madsen, H.R., 2015. CD107a as a marker of activation in chicken cytotoxic T cells. J. Immunol. Methods 419, 35–47. doi:10.1016/j.jim.2015.02.011

Wynn, R.F., Hart, C.A., Corradi-Perini, C., O’Neill, L., Evans, C.A., Wraith, J.E., Fairbairn, L.J., Bellantuono, I., 2004. A small proportion of mesenchymal stem cells strongly expresses functionally active CXCR4 receptor capable of promoting migration to bone marrow. Blood 104, 2643–2645. doi:10.1182/blood-2004-02-0526

Yang, J.-X., Zhang, N., Wang, H.-W., Gao, P., Yang, Q.-P., Wen, Q.-P., 2015. CXCR4 receptor overexpression in mesenchymal stem cells facilitates treatment of acute lung injury in rats. J Biol Chem 290, 1994–2006. doi:10.1074/jbc.M114.605063

York, V.A., Milush, J.M., 2015. Ex vivoHuman Natural Killer (NK) Cell Stimulation and Intracellular IFNγ and CD107a Cytokine Staining. Bio Protoc 5.

Yu, J.M., Wu, X., Gimble, J.M., Guan, X., Freitas, M.A., Bunnell, B.A., 2011. Age-related changes in mesenchymal stem cells derived from rhesus macaque bone marrow. Aging Cell 10, 66–79. doi:10.1111/j.1474-9726.2010.00646.x

Yue, R., Zhou, B.O., Shimada, I.S., Zhao, Z., Morrison, S.J., 2016. Leptin Receptor Promotes Adipogenesis and Reduces Osteogenesis by Regulating Mesenchymal Stromal Cells in Adult Bone Marrow. 18, 782–796. doi:10.1016/j.stem.2016.02.015

Zhou, B.O., Yue, R., Murphy, M.M., Peyer, J.G., Morrison, S.J., 2014. Leptin-receptor-expressing mesenchymal stromal cells represent the main source of bone formed by adult bone marrow. 15, 154–168. doi:10.1016/j.stem.2014.06.008

Zhou, Y., Day, A., Haykal, S., Keating, A., Waddell, T.K., 2013. Mesenchymal stromal cells augment CD4+ and CD8+ T-cell proliferation through a CCL2 pathway. Cytotherapy 15, 1195–1207. doi:10.1016/j.jcyt.2013.05.009

